# GABA, glutamatergic dynamics and BOLD contrast concurrently assessed using functional MR spectroscopy during a cognitive task

**DOI:** 10.1101/2023.05.02.539017

**Authors:** Alexander R. Craven, Gerard Dwyer, Lars Ersland, Katarzyna Kazimierczak, Ralph Noeske, Lydia Brunvoll Sandøy, Erik Johnsen, Kenneth Hugdahl

## Abstract

A recurring issue in functional neuroimaging is how to link task-driven hemodynamic BOLD-fMRI responses to underlying neurochemistry at the synaptic level. Glutamate and GABA, the major excitatory and inhibitory neurotransmitters respectively, are typically measured with MR spectroscopy (MRS) sequences in a resting state, in the absence of a task. We propose to resolve this disconnect by applying a novel method to concurrently acquire BOLD, Glx and GABA measurements from a single voxel, using a locally adapted MEGA-PRESS sequence implementation which incorporates unsuppressed water reference signals at a regular interval, allowing continuous assessment of BOLD-related linewidth variation for fMRS applications.

Healthy subjects (N = 81) performed a cognitive task (Eriksen Flanker) which was presented visually in a task-OFF, task-ON block design, with individual event stimulus timing varied with respect to the MRS readout. BOLD data acquired with the adapted MEGA-PRESS sequence were correlated with data acquired using a standard fMRI EPI sequence as a means of validating the concurrent approach: a significant (although moderate) correlation was observed, specific to the fMRS-targeted region of interest. We additionally present a novel linear model for extracting modelled spectra associated with discrete functional stimuli, building on well-established processing and quantification tools. Behavioural outcomes from the Flanker task, and activation patterns from the BOLD-fMRI sequence, were as expected from the literature. fMRS-assessed BOLD response correlated strongly with slowing of response time in the incongruent Flanker condition. Moreover, there was a significant increase in measured Glx levels (~8.8%), between task-OFF and task-ON periods. These findings verify the efficacy of our functional task and analysis pipelines for the simultaneous assessment of BOLD and metabolite fluctuations in a single voxel. As well as providing a robust basis for further work using these techniques, we also identify a number of clear directions for further refinement in future studies.

**Highlights:**

- Concurrent measurement of temporally resolved metabolite estimates and local BOLD-related signal changes is demonstrated
- In-vivo, GABA-edited functional ^1^H-MRS data were collected from 81 healthy subjects whilst they performed a cognitive task
- Robust task-related increases in measured Glutamate+Glutamine were observed in the Anterior Cingulate Cortex
- Moderate correlation was observed between BOLD contrast strength as assessed by fMRS and fMRI techniques, specific to the prescribed region
- A novel technique for extraction of block- or event-related sub-spectra is demonstrated

**Graphical Abstract:** A novel sequence adaption for concurrent measurement of GABA-edited spectroscopy and functional BOLD changes is demonstrated on N=81 healthy subjects. Significant changes in measured Glutamate+Glutamine concentration are found, along with the expected behavioural and BOLD effects in response to a Flanker task.

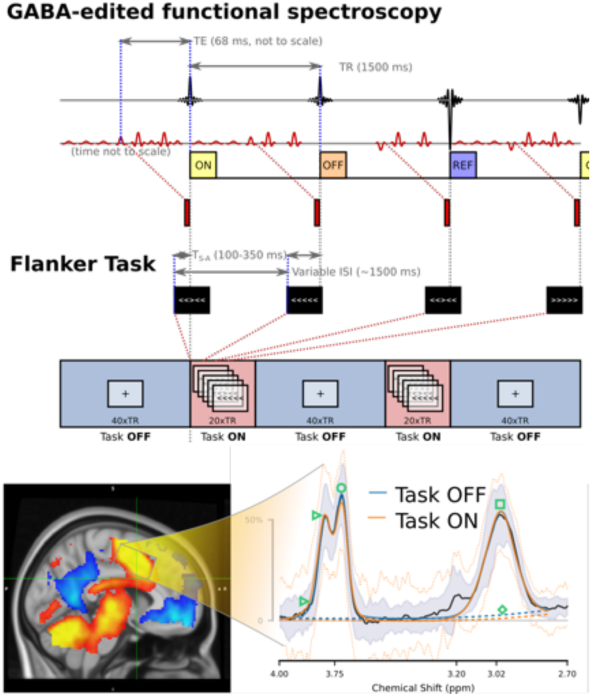

## 1 Introduction

Since the advent of blood oxygen level dependent (BOLD) functional magnetic resonance imaging (fMRI) techniques for advanced neuroimaging, such techniques have been widely adopted for studying patterns of activation in neural systems and networks of the healthy and the diseased brain (see ^1,2^ for overviews). While informative, a more comprehensive understanding of complex mental phenomena requires consideration across multiple complementary “levels of explanation” ^3^. Of particular interest is harmonising observed findings with putative changes on the cellular (neurotransmitter) level, using magnetic resonance spectroscopy (MRS) techniques. Such techniques allow non-invasive, quantitative assessment of neurotransmitter concentrations in the living brain, including measurement of glutamate (Glu) and γ-aminobutyric acid (GABA), the primary excitatory and inhibitory neurotransmitters respectively ^4^. Typically, MRS data would be collected in a separate acquisition run within the same scanning session, but minutes before or after the BOLD-fMRI sequence, which is not informative of the change in neurochemistry taking place at the time of the mental event. What complicates matters further is that the MRS data are typically acquired during a resting condition, irrespective of whether the BOLD data are acquired during resting or during task performance. This absence of direct concurrency makes it difficult to draw firm conclusions regarding the instantaneous, dynamic inter-relation of the measured signals. Thus, there is a need to develop new approaches for simultaneous acquisition of MRS and BOLD data to overcome the problem of separate time scales. This would be a priority challenge when it comes to relating brain activity to mental events in the sense of cognitive performance, since all markers of BOLD activation during a cognitive task would be gone long before the MRS sequence is applied.

There are however several challenges associated with MRS techniques, particularly when considered in relation to task performance. Firstly, the metabolite signals of interest are several orders of magnitude weaker than the water signal which forms the basis of BOLD-fMRI measurements. This means that for reliable measurement, the MRS signal must be acquired from a fairly large volume, over a large number of repetitions (hence, a long overall scan time) – typically volumes in the order of 20 mL and scan lengths in the 5-10 minute range may be needed. These are in direct competition with demands for good anatomical specificity and high temporal resolution, as necessary for observing subtle, short-term functional dynamics localised to a particular brain region or network. Moreover, while each of the MRS-observable metabolites exhibits a unique, characteristic spectrum by which it may be identified, there is often substantial overlap between these – to the point where some metabolites are indistinguishable at common clinical field strengths, while others require specialised acquisition techniques to resolve them from stronger signals in the vicinity which would otherwise effectively bury the signal of interest.

GABA is one such signal: at typical in-vivo concentrations, the 3 ppm GABA peak is usually engulfed by much stronger signals from Creatine (Cr) in the same part of the spectrum, while the 1.89 ppm peak is often masked by the large 2.0 ppm N-acetylaspartate signal (NAA). One approach to isolating the GABA signal is to use selective editing techniques ^5,6^, such as the commonly-used MEGA-PRESS sequence. In such a sequence, a sub-spectrum in which coupling to GABA spins around 3 ppm has been selectively refocussed with a so-called “editing” pulse is subtracted from a second sub-spectrum in which it has not. Taking the difference between these edit-ON and edit-OFF sub-spectra yields a difference spectrum (DIFF) containing only signals which have been impacted by the selective refocussing (editing) pulse, i.e. GABA around 3 ppm and a handful of co-edited metabolite and macromolecule signals.

Such a sequence places additional demands on the amount of signal which must be acquired to achieve an acceptable Signal-to-Noise Ratio (SNR). This is due to the need for two distinct yet well matched sub-spectra and losses associated with imperfect editing, compounding already weak signal due to comparatively low biological concentrations of GABA. Accurate frequency alignment of individual transients is also critical to avoid artefacts associated with imperfect subtraction of the sub-spectra.

Although ostensibly selective, the editing pulse in a typical MEGA-PRESS sequence necessarily has a finite bandwidth and thus will co-edit certain other metabolites having spins in nearby spectral regions. This gives rise to a distinct double peak associated with the C2 multiplets of Glutamate and Glutamine around 3.75 ppm in the difference spectrum, and a strong negative peak associated with NAA around 2.0 ppm. It also gives rise to some additional signal underlying the GABA signal of interest at 3 ppm, tentatively attributed to homocarnosine and lysine-containing macromolecules and denoted herein as MM3co. This co-edited signal contributes roughly 50% to the measurable signal in that part of the spectrum ^7^, depending on experimental conditions. As reliable separation of these remains a challenge, GABA estimates obtained in this way are usually reported as GABA+: GABA plus contribution from co-edited signals in the area.

Further to the competing demands of high SNR, good temporal resolution and reasonable anatomical specificity, MRS measured in the presence of a cognitive task may be confounded by BOLD-related changes in signal relaxation and local shim quality. The effective transverse relaxation time (T_2_*) is inversely proportional to spectral linewidth, which may affect the separability of certain metabolites or their unambiguous extraction from background signals in the acquired spectra, hence may potentially introduce biases in concentration estimates obtained by certain algorithms. Reliable consideration of MRS data acquired under functional (task) conditions is contingent on accurate characterisation and consideration of such changes.

To address some of these challenges, we applied a novel, locally adapted MEGA-PRESS implementation in the present study, which incorporates unsuppressed water reference signals at a regular interval within the edited acquisition scheme. This would allow quantification of the BOLD effect concurrently throughout the MRS acquisition period and permit improved dynamic linewidth assessment. A well-known and reasonably demanding cognitive task was used (based on the Eriksen Flanker task ^8^), coupled with a novel linear decomposition procedure to extract reconstructed sub-spectra corresponding with functional events or blocks of interest. This allows the efficacy of these techniques to be assessed.

While the present report focusses on fMRS outcomes for healthy subjects, these data were collected as part of a larger study also including a clinical cohort. The current report therefore represents a validation of the underlying methods, as a precursor to forthcoming analyses combining the methods described herein with additional imaging modalities and in-depth behavioural investigation, additionally incorporating clinical subjects.

## 2 Methods

### 2.1 In-vivo data collection

#### 2.1.1 Subject recruitment and demographics

Healthy subjects (N = 81) were recruited from the Bergen local community and at Haukeland University Hospital through posters and through an article in a local newspaper (Bergens Tidende) covering the Bergen City and Vestland County regions in Western Norway. There were 44 males and 37 females, with mean age of 34.37 ± 11.2 years, range 19-62 years. All potential subjects were screened before inclusion in the study regarding previous history of major head injuries, medical implanted devices, substance abuse, neurological- and medical illnesses. Written informed consent was acquired from all subjects prior to the study. Subjects were instructed not to consume caffeinated drinks or to use substances containing nicotine in the two hours before coming to the MR scanner. The study was approved by the Regional Committee for Medical Research Ethics in Western Norway (REK Vest # 2016/800)

#### 2.1.2 MR scanning protocol

The scanning protocol included a high-resolution T_1_ structural acquisition: fast spoiled gradient (FSPGR) sequence, 188 sagittal slices, 256×256 isometric 1 mm voxels, 12 degree flip angle, TE/TR approximately 2.95, 6.8 ms respectively. Acquisition of functional MRS data followed: GABA-edited spectroscopy data were collected with a locally-modified sequence detailed in section 2.2, whilst subjects performed the functional task described in section 2.3. Subsequently, BOLD fMRI data were collected as subjects performed a similar functional task, with an echo-planar imaging (EPI) sequence, TE=30 ms, TR=2500 ms, 90 degree flip angle, 36 slices of 128 x 128 voxels (1.72×1.72 mm), 3.0 mm slice thickness with 0.5 mm gap (3.5 mm slice spacing); 240 volumes for a total acquisition time of 600 s. Resting state fMRI (rs-fMRI) data were also acquired (480 seconds, 30 slices, TR=2000 ms, otherwise similar to the task fMRI sequence), recorded along with photoplethysmographic (PPG) pulse data; these resting state data are not considered in the present analysis. Note that gradient-heavy fMRI acquisitions were performed *after* the MR spectroscopy, to minimise the impact of thermal drift ^9^ on the edited fMRS acquisition; this precluded counterbalancing of the fMRI and fMRS acquisition order.

### 2.2 MEGA-PRESS sequence adaption

To facilitate simultaneous acquisition of time-resolved MRS data and robust assessment of BOLD signal changes in response to the cognitive task, local extensions to the MEGA-PRESS sequence were adopted. These build upon the standard GE MEGA-PRESS implementation, adding per-TR trigger pulses (500 µs after the end of the excitation RF pulse) to allow for precise synchronisation with an external stimulation paradigm. Moreover, the local sequence implementation has the option to periodically disable the CHESS water suppression pulses, at a user-specified interval – thereby periodically acquiring a water-unsuppressed reference signal within the regular GABA-editing sequence.

GABA-edited data were acquired from a 22×36×23 mm (18.2 mL) voxel placed medially in the Anterior Cingulate Cortex (ACC) (see section 2.2.1), with TR=1500 ms (note discussion in Section 4.3), TE=68 ms and with 15 ms sinc-weighted Gaussian editing pulses at 1.9/7.46 ppm for edit-ON/-OFF respectively; 2-step phase cycling, 700 transients alternating edit-ON/-OFF (total scan time just under 18 minutes) and with CHESS suppression pulses disabled in every third transient; the resultant pulse sequence timing is illustrated in Figure 1. These parameters ensured that the sequence remained balanced with respect to the number of edit-ON, edit-OFF and water-unsuppressed reference (WREF) signals acquired at each phase-cycle step, and with respect to any carryover impact of residual, unrelaxed signals (particularly from the water-unsuppressed acquisitions) on subsequent transients, with an equal proportion of edit-OFF and edit-ON sub-spectra at each phase cycle step being preceded by a water-unsuppressed reference acquisition.

**Figure 1:**
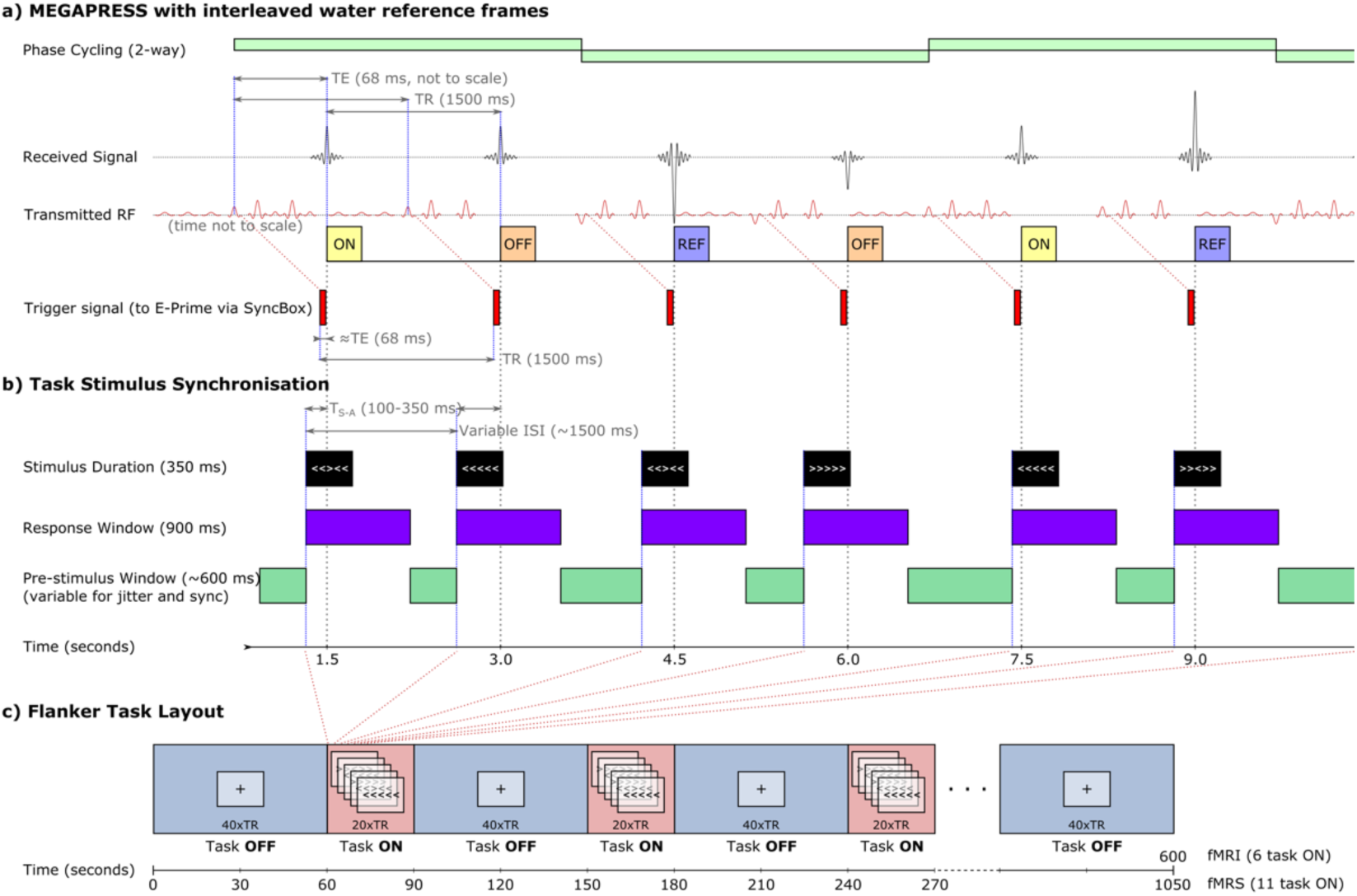
Timing of the fMRS acquisition (a), synchronisation to task stimuli (b) and structure of the associated Flanker task (c)

A summary of key sequence and hardware parameters is presented in an MRS-in-MRS ^10^ checklist, in the supplementary material.

#### 2.2.1 Voxel placement and early piloting

Meaningful positioning of the volume of interest (VOI) can be a challenge for single-voxel fMRS acquisitions. To determine a suitable positioning scheme for the present study, a small pilot study was performed where fMRI data were collected from 10 pilot subjects, whilst they performed the same Flanker task as used in the main study. fMRI block analysis was performed on these pilot data, using a similar procedure to that described in the main analysis (see section 2.5). A local script implemented in Matlab [v2021a; MathWorks Inc., Natick, MA, USA] was used to iteratively optimize the position, dimensions and orientation of a cuboid VOI, wherein voxels intersecting the group-mean Z-maps for the task-ON functional response, in Gray or White matter in the ACC were prioritised, and intersection with task-OFF activation, overlap with the Precuneus structure and probable CerebroSpinal Fluid (CSF) or non-brain content were penalised; for this purpose, the standard Tissue Probability Map (TPM) ^11–13^ shipped with the SPM12 software (https://www.fil.ion.ucl.ac.uk/spm/software/spm12/) was used, with anatomical structures defined from the Harvard-Oxford cortical atlas^14^ shipped with the FSL software (FMRIB’s Software library, http://www.fmrib.ox.ac.uk/fsl).

With a target volume around 19 mL and side lengths constrained to 27+/-5 mm, position, dimensions and angulation were iteratively refined to converge on an optimal placement (at group level) covering the major functional activation in the ACC during task-ON blocks, while avoiding problematic regions and nearby brain regions (particularly the precuneus) expected to exhibit differential activation patterns (hence, potentially confound the results) during task-OFF blocks. This “optimal” placement was used to guide individual placements during the main study. The resulting placements achieved for the main study are presented in Figure 2.

**Figure 2:**
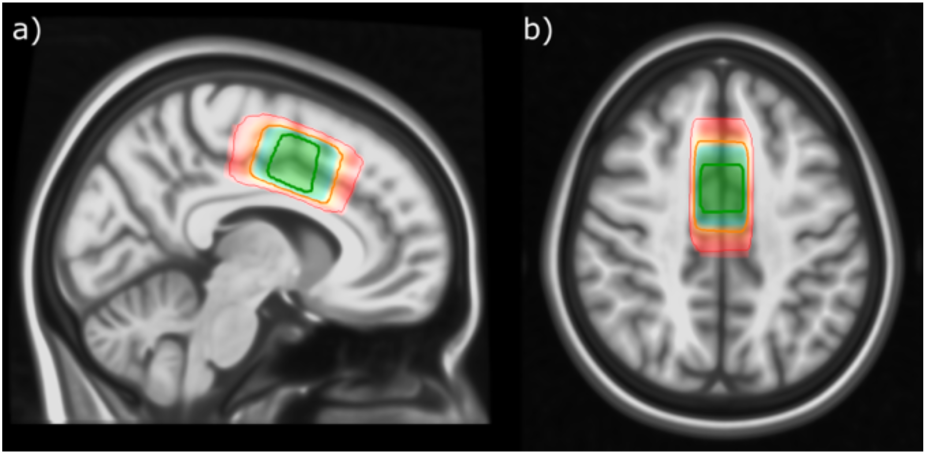
Placement of the fMRS VOI for the healthy subjects, mapped to standard space. Contours indicate [5,50,95]-percentile coverage of the achieved placement across subjects

### 2.3 Functional paradigm – Flanker task

A variation of the Erikson Flanker task^8^ was used to drive cognitive load. Tasks of this nature have well-documented behavioural and brain functional characteristics both in healthy subjects and in certain clinical groups, including in patients with schizophrenia ^15–18^. A Flanker task using visually presented arrows was chosen so as to remain largely independent of auditory and language pathways which may be aberrant in certain conditions of interest, or may be confounded by the noisy scanner environment. In this task, a set of arrow glyphs are presented in each trial, with a central “target” arrow surrounded by four “flankers”. The task of the subject is to indicate the direction of the central target arrow by pressing a response button. If the direction of the target and the flankers match (for example, “> > > > >” or “< < < < <”), the trial is “congruent”. If the flankers point in the opposite direction to the target (“< < > < <” or “> > < > >”), this is a cognitively more demanding “incongruent” trial. The present implementation did not incorporate stimuli with “neutral” flankers, and always presented flanker and target stimuli concurrently (stimulus onset asynchrony SOA=0).

The paradigm was implemented in E-Prime 2.0 SP1 [2.0.10.353: Psychology Software Tools Inc., Pittsburgh, PA, https://pstnet.com/], and presented from a separate PC adjacent to the MR system console. The arrow stimuli were presented during task-ON blocks through a set of goggles mounted on the head coil [NordicNeuroLab AS (NNL), Bergen, Norway, http://nordicneurolab.com/, note declaration of interest] in light grey on a black background, with relatively small font and spacing to ensure that the stimulus remained near the parafoveal field of view; see Supplementary Figure 1d. A central fixation cross (Supplementary Figure 1c) was presented between each trial, and during OFF blocks. The paradigm was run as a standard block-event design, beginning with a 60-second task-OFF block followed by alternating 30-second task-ON blocks and 60-second task-OFF blocks. The 18-minute fMRS acquisition allowed for 11 task-ON blocks, while the 10-minute fMRI acquisition accommodated six blocks. This is summarised in Figure 1c. Within each task-ON block, one trial was presented per TR (i.e., one every 1500 ms); this timing allowed a total of 220 stimuli for the fMRS acquisition, and 120 stimuli for the shorter fMRI acquisition – of which a randomly-selected 40% were incongruent and the remainder were congruent. Trial onset timing was jittered randomly such that stimuli were presented T_S-A_ = 100-350 ms before the MRS trigger ^19^; stimuli for each trial were presented for 350 ms, with a nominal 800 ms response window (measured from the beginning of presentation). As such, the nominal inter-stimulus interval (ISI) was 1500 ms, but varied somewhat on a per-trial basis according to the jittered onsets with respect to the fixed periodic TR. The subjects held a set of response grips (NNL), with which they were instructed to respond promptly to indicate the direction of the central “target” arrow – pressing with the left index finger when the central arrow points to the left, or with the right when it points to the right.

Trigger signals from the scanner were received by E-Prime via an NNL SyncBox, with synchronisation between the scanner and the paradigm updated continuously. Since stimulus onset preceded receipt of triggers from the scanner, synchronisation was achieved by way of a feedback loop, using the difference between nominal and achieved intervals to iteratively tune compensatory adjustment factors – as such, actual intervals deviated slightly from nominal; details of the actual achieved timing for stimulus presentation, MR trigger receipt and the subjects’ response were recorded and used in all subsequent analysis. In the case of the fMRI acquisition, where the TR of 2500 ms did not always align to the ISI of around 1500 ms, triggers were only issued and adjusted to at the beginning of each new task-ON or task-OFF block.

Subjects were instructed on the nature of the task before the scanning session commenced. This included presentation of sample congruent and incongruent stimuli (in printed form) to ensure that they understood the task, and familiarisation with the response grips. Further instruction was presented through the goggles immediately before the fMRS task, as the subject lay in the scanner. This incorporated feedback from the subject, thereby validating the communication between key components for each session: the stimulation PC, visual system, human subject, response grips and the scanner’s trigger signals. Instruction was provided in Norwegian or English depending on each subject’s preference; sample instruction screens are presented in Supplementary Figure 1a,b.

Individual response data were processed using in-house tools (see section 2.6). Missing responses (where no button press was registered within the response window) were flagged, as were invalid responses, such as cases where more than one button press was registered in the response window. After excluding missing and invalid responses, basic task performance metrics of mean Reaction Time (RT) and % Response Accuracy (RA) were evaluated. This was done separately for each subject, each functional scan (fMRS and subsequent fMRI), and for each flanker condition (congruent vs incongruent). Median and median absolute deviation (MAD) across subjects for each parameter were subsequently evaluated, as reported in section 3.1. Outlier-resistant methods were chosen to minimise the risk of a handful of underperforming subjects driving any eventual outcomes. Achieved ISI and proportion of congruent vs incongruent stimuli were also assessed to verify correct execution of the paradigm itself.

### 2.4 fMRS data processing and quantification

Functional MEGA-PRESS data were initially loaded using the GannetLoad function from Gannet version 3.1 ^20^. While this function incorporates a comprehensive processing pipeline, only part of this functionality was used in the present study. After import, eddy-current correction ^21,22^ and global zero-order phase correction using a dual-Lorentzian model for Cr and choline (Cho) were performed. Robust Spectral Registration ^23^ was subsequently applied independently for edit-OFF and edit-ON transients, and for each phase cycling step, before alignment of the sub-spectra by spectral registration ^24^ – all using basic Gannet functionality. Individual transients were taken for further processing.

#### 2.4.1 Subject Motion and Discontinuities

Inevitably with such long acquisitions as performed here, subject motion must be considered. As the subject moves, not only will the measurement be acquired from a slightly different region from that initially prescribed, but also the local shim conditions change – the latter leading to abrupt changes in the spectral line shape. Such motion may often be observed as an abrupt change in the estimated water frequency.

A coarse estimate of water frequency was obtained from the location of the peak voxel near the expected location of (residual) water in each transient, after Fourier transformation (FFT) and zero-fill. Abrupt changes in the water frequency were interpreted as indicative of changing scanning conditions – most likely as a result of subject motion. Such changes were identified from the assessed water frequency after applying a four-point median filter, using Matlab’s `findchangepts` function, operating in linear mode to detect abrupt changes in mean and slope. A minimum distance (`MinDistance`) between changepoints was set at 8 transients. After a series of assays with different thresholding criteria, a threshold on the minimum improvement in total residual error (`MinThreshold`) was set at eight times the variance of the signal, after subtraction of a third-order polynomial fit to remove variance associated with gradual drift (e.g., thermal drift); this threshold was then clipped to a lower limit of 3×10^−5^ to prevent over-characterising more stable acquisitions. Data were partitioned according to the identified change points, as illustrated in Figure 3b.

**Figure 3.**
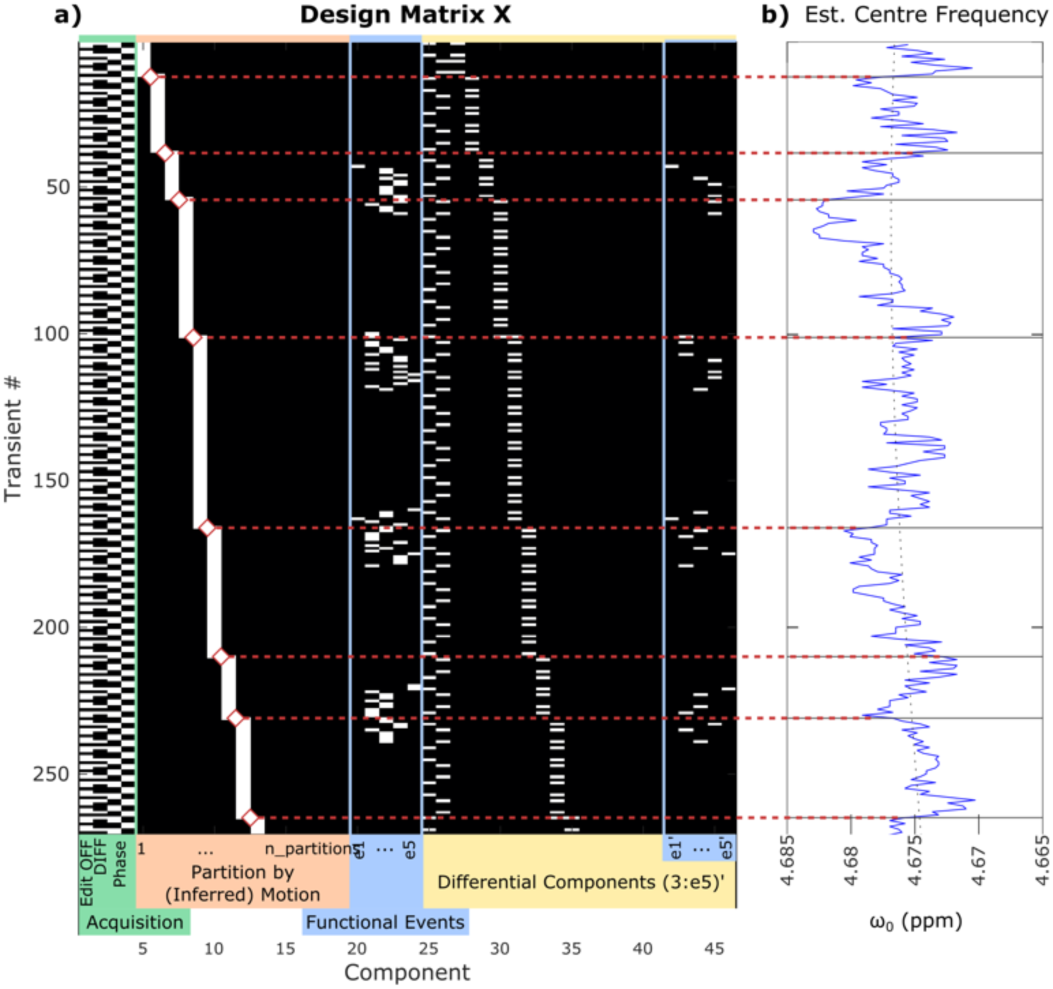
Sample design matrix (a) and partitioning according to motion inferred from the water frequency (b), for the first three minutes of data from a single representative subject.

#### 2.4.2 Metabolite sub-spectra: a linear model for spectral combination

A linear model of the form Y=XB+U was constructed, wherein each acquired transient (Y_n,*_) is expressed as a combination of a normal, baseline edit-OFF spectrum (X_*,1_), with periodic contributions associated with editing (the baseline DIFF spectrum, X_*,2_), periodic covariate associated with each of the two phase-cycle steps (X_*,3:4_), and additional covariates partitioning the data according to inferred motion (X_*,4+(1:n_partitions)_).

Functional events were binned according to the achieved interval between stimulus and acquisition (where the latter is defined by the receipt of the trigger pulse from the scanner, 500 µs after the end and 2344 µs after the centre of the 90 degree RF pulse). Five bins were defined, with edges at [100, 183, 267, 350] ms, open at either end, lower limits inclusive; the inner three bins evenly cover the nominated 100-350 ms T_S-A_ range. This resulted in approximately 48 task-ON metabolite transients per bin (with some individual variation), and 320 task-OFF metabolite transients. These bins (bin_1-5_) were incorporated into the design matrix as columns X_*,e1:e5_. Note that columns (3:e5) capture variability common to the edit-OFF and DIFF (sub-)spectra; these columns were subsequently replicated and masked to capture differential changes associated with the respective variables; corresponding differential masked event columns are designated X_*,(e’1:e’5)_. A representative design matrix for a single subject is presented in Figure 3a. Two further variations of this design matrix were used, to extract sub-spectra corresponding with different functional stimuli, and with different periods within the task-ON block. For the former, the event-related columns X_*,(e1:e4)_ instead selected transients according to the stimulus and response: congruent accurate, congruent inaccurate, incongruent accurate, incongruent inaccurate. Approximately 88 and 59 task-ON metabolite transients were available for the congruent and incongruent components respectively, with subdivision according to accuracy dependent on individual performance. For subdivision of the task-ON blocks, event-related columns X_*,(e1:e5)_ selected successive 7.5-second portions of all task-ON blocks ([0-7.5, 7.5-15, 15-22.5, 22.5-30] s), and the first part of the following task-OFF block (30-37.5 s).

The system was solved stepwise: an initial fit was performed using only the components of interest (edit-OFF, DIFF and masked event bins), with the covariate components subsequently modelled to the residuals from that fitting. In both cases the Matlab `decomposition` function was used for complete orthogonal decomposition to yield a minimum norm least-squares solution. Removal of “bad” transients (such as those corrupted by motion) was performed based on the residual error after fitting, where the Z-score of the residual error for a given transient exceeded 2.5. After removal of bad transients, the two fitting steps were repeated until the relative improvement in `norm_residuals` from the previous step was below 10^−8^. Baseline edit-OFF and DIFF spectra were extracted directly from the corresponding components X_*,(1,2)_, while edit-OFF and DIFF components binned according to functional events (bin_1-5_) were evaluated arithmetically from components X_*,(e1:e5)_ and X_*,1_, and X_*,(e’1:e’5)_ and X_*,2_, respectively. Linewidth matching of the extracted spectra was performed with a Lorentzian filter (exponential decay in the time domain), using Cho and Cr from the associated edit-OFF sub-spectra to assess linewidth; line broadening was performed to match the maximum measured linewidth across all extracted spectra for the individual subjects, capped at 130% of the median to avoid matching to strong outlier targets. Finally, singular value decomposition (SVD) was used to remove residual water signal. Extracted spectra were then fit using Gannet’s `GABAGlx` model, which fits GABA+ with a Gaussian peak around 3.02 ppm and Glx as a pair of Gaussian peaks around 3.71 and 3.79 ppm. Water-scaled estimates with tissue-class correction ^25^ are reported.

#### 2.4.3 Quality Control

Three rejection criteria were applied to the extracted DIFF spectra, in series:

- R1 rejects fits where the full-width at half maximum (FWHM) linewidth of the fitted NAA or GABA+ peak from the extracted DIFF spectrum exceeded 12 or 30 Hz, respectively.
- R2 rejects fits where the SNR of the fitted NAA peak is extraordinarily low, below 20 (noting that quite low SNR may be expected in some of the extracted event-related spectra)
- R3 rejects extreme outliers: GABA+ or Glx estimates differing from the median by more than five times the Median Absolute Deviation (MAD) across all spectra surviving R1 and R2.

#### 2.4.4 Unsuppressed water signal: T_2_* and BOLD assessment

For each timepoint (TR), an unsuppressed water-reference spectrum WREF was modelled as the weighted average of temporally near water-unsuppressed transients – weighted with a gaussian kernel of standard deviation 1.5 seconds. For each such timepoint, this modelled water signal was fit with a pseudo-Voigt function superimposed on a linear baseline, with centre frequency, amplitude above baseline, zero-order phase and FWHM linewidth logged – the latter being closely tied to T_2_*. These data were subsequently processed to identify abrupt changes in the signal, as may arise from subject motion. Outliers in the FWHM fit were flagged, where the deviation from the mean exceeded four times the MAD after subtracting a third-order polynomial fit to remove any broad effects (such as those related to thermal drift). Discontinuities in the frequency, amplitude or zero-order phase were also identified using linearly penalized segmentation ^26^ with a kernelized mean change cost function ^27,28^, implemented in the ruptures toolkit ^29^.

A linear model, similar to that described for the metabolite signal in section 2.4.2, was constructed. Unit impulses at the recorded functional event onsets were convolved with a dual-gamma haemodynamic response function (HRF) model to create a model for the expected BOLD signal, (X_*,1_), with covariates for phase cycling step (X_*,2:3_), and inferred subject motion (X_*,3+(1:n_partitions)_). The model was fit with linear least-squares regression, with the BOLD model coefficient Y_1_ (roughly equivalent to ΔFWHM_water_, the overall change in water linewidth) reflecting the strength of the individual’s BOLD response as assessed from WREF signals during the fMRS acquisition.

### 2.5 fMRI data processing

fMRI block analysis was performed using FEAT (FMRI Expert Analysis Tool version 6.00, part of FSL). Processing included motion correction using MCFLIRT ^30^, masking of non-brain data using BET ^31^, spatial smoothing (5 mm FWHM Gaussian kernel), and linear registration to a high resolution standard space structural template using FLIRT ^30,32^, subsequently refined with non-linear registration using FNIRT ^33,34^. The MNI-512 (non-linear, 6^th^ generation) template ^35^ was used as a registration target. Time-series statistical analysis was performed using FILM with local autocorrelation correction ^36^; the resulting statistical images in subject space were thresholded non-parametrically ^37^ using clusters determined by Z>3.1 and a corrected cluster significance threshold of p_cluster_=0.05. Individual registration outcomes were subject to visual inspection, and higher level (group average) statistics evaluated after transformation into standard space; group average response allowed visual inspection of the task response and selection of control VOIs, but was not subject to further quantitative analysis.

VOIs were defined using the individually prescribed fMRS voxel geometry in the anterior cingulate cortex (VOI_fMRS,ACC_). Based on the group average fMRI response in standard space, additional VOIs with similar volume to the fMRS acquisition (around 18.2 mL) were defined in the temporal pole (VOI_temporal_ around [−38.7, 10.6, −33.6] mm), the precuneus near the posterior cingulate (VOI_precuneus_ around [0.5, −56, 20] mm), the occipital (VOI_occipital_ around [2.83, −93, 12.3] mm) and the anterior cingulate cortex (VOI_ACC_ around [0.5, 3.5, 45.5] mm) in standard MNI 512 template space, subsequently transformed to the individual subject geometry. Median Z-score across each VOI was extracted, without thresholding. Subjects where the median Z-score within the spectroscopy voxel was not positive were considered non-conformant: either the individual’s task performance did not elicit the expected functional response, or the achieved positioning of the fMRS voxel did not demonstrably capture this response; see Figure 6a.

### 2.6 Numerical and Statistical Analysis

Outcomes of the behavioural, fMRI and fMRS tasks were collated and analysed using locally developed scripts implemented in Python (v3.7.3), with data analysis using methods from pandas ^38^ (v1.5.2) and NumPy ^39^ (v1.23.5). Numeric and statistical methods were derived from the SciPy ^40^ (v1.9.3), pingouin ^41^ (v0.5.2) and statsmodels ^42^ (v0.13.5) libraries. Visualisation was built on tools from matplotlib ^43^ (v3.3.4), seaborn ^44^ (v0.11.2) and statannotations ^45^ (v0.5).

Where appropriate, compatibility of variance was assessed using the Fligner-Killeen method ^46^, and normality was assessed with the Shapiro-Wilk method ^47^. In cases where Shapiro-Wilk indicated non-normality, the Wilcoxon Signed Rank test ^48^ was used for hypothesis testing between related samples. Confidence intervals derived from parametric bootstrapping (10,000 permutations) are denoted CI_boot,95%_. In cases where adjustment for multiple comparisons was performed, Holm-Bonferroni ^49,50^ correction was applied; adjusted p-values are denoted p_holm_, with a corrected significance threshold defined as p_holm_<0.05; uncorrected p-values are denoted p_unc._. Where correlation coefficients are reported, these are robust Spearman correlations using the skipped method ^51,52^ to exclude bivariate outliers.

## 3 Results

### 3.1 Behavioural Outcomes

Basic behavioural outcomes from the Flanker task are summarised in Table 1 and Figure 4. Shapiro-Wilk testing indicated that most metrics followed a non-normal distribution – in particular, those measures relating to accuracy presented a highly skewed distribution. This motivated the adoption of the Wilcoxon Signed Rank test for hypothesis testing. Strongly significant differences (p_holm_<0.001) were seen between congruent and incongruent trials for the RA, RT and RA/RT parameters, during both the fMRS and fMRI task-ON blocks. Comparing the fMRS and subsequent fMRI task performance, increases in RA and RA/RT were observed in the latter, which was significant (p_holm_<0.01) for incongruent trials.

**Figure 4.**
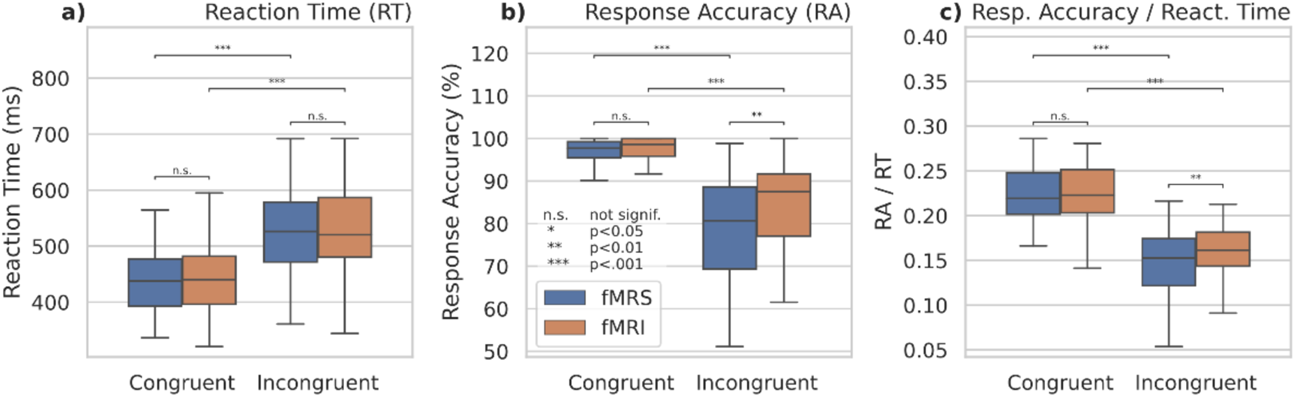
Outcomes from the behavioural task

**Table 1.**
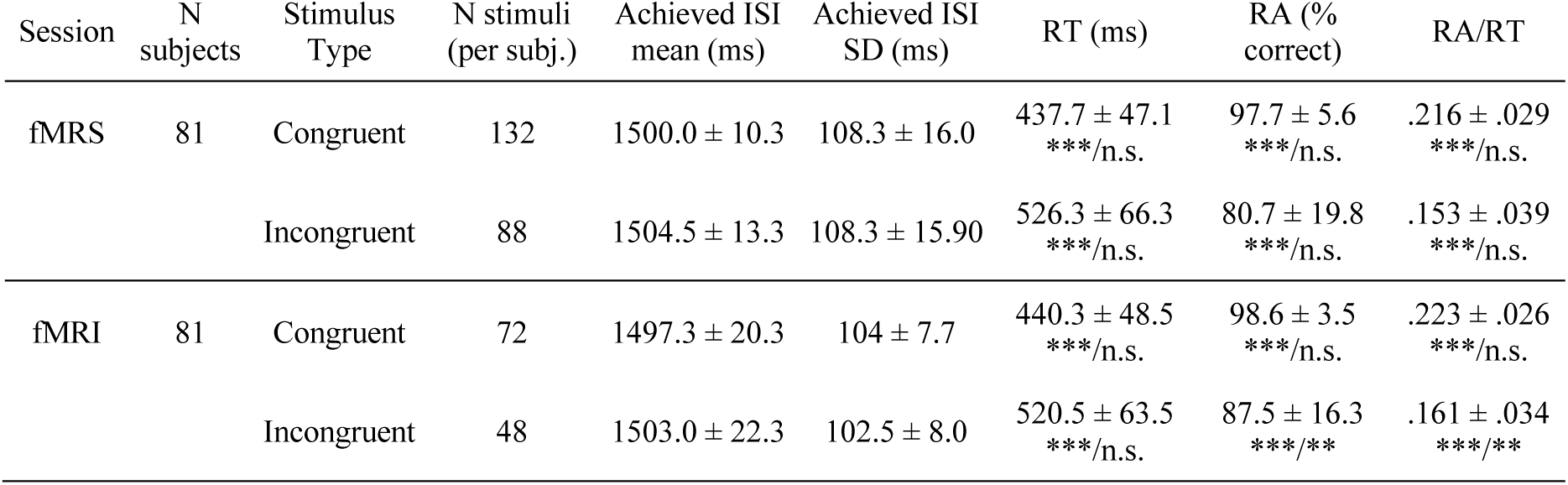
Behavioural outcomes from the Flanker task; values are quoted as Median +/- Median Absolute Deviation (MAD) of per-subject outcomes. Significant differences are indicated across stimulus type and session (denoted type/session, *** p_holm_<0.001, ** p_holm_<.01, * p_holm_<.05, n.s. not significant). ISI: Inter-stimulus interval, RA: Response Accuracy, RT: Response Time

| Session | N subjects | Stimulus Type | N stimuli (per subj.) | Achieved ISI mean (ms) | Achieved ISI SD (ms) | RT (ms) | RA (% correct) | RA/RT |
| --- | --- | --- | --- | --- | --- | --- | --- | --- |
| fMRS | 81 | Congruent | 132 | 1500.0 ± 10.3 | 108.3 ± 16.0 | 437.7 ± 47.1<br>*** / n.s. | 97.7 ± 5.6<br>*** / n.s. | .216 ± .029<br>*** / n.s. |
|  |  | Incongruent | 88 | 1504.5 ± 13.3 | 108.3 ± 15.90 | 526.3 ± 66.3<br>*** / n.s. | 80.7 ± 19.8<br>*** / n.s. | .153 ± .039<br>*** / n.s. |
| fMRI | 81 | Congruent | 72 | 1497.3 ± 20.3 | 104 ± 7.7 | 440.3 ± 48.5<br>*** / n.s. | 98.6 ± 3.5<br>*** / n.s. | .223 ± .026<br>*** / n.s. |
|  |  | Incongruent | 48 | 1503.0 ± 22.3 | 102.5 ± 8.0 | 520.5 ± 63.5<br>*** / n.s. | 87.5 ± 16.3<br>*** / ** | .161 ± .034<br>*** / ** |

### 3.2 Functional Outcomes

#### 3.2.1 BOLD assessment by fMRI and fMRS

Group-average spatial response for the fMRI task is presented in Figure 5, showing strong task-related activations in fronto-temporo-parietal regions and the ACC/SMA as would be expected for a cognitively demanding task such as the Flanker task, with parietal/precuneus and ventromedial inferior frontal regions more activated during task-OFF periods between the task-ON blocks. Cluster statistics, location of local maxima, and corresponding atlas structures are presented in Supplementary Section C, with additional slices shown in Supplementary Figure 2.

**Figure 5.**
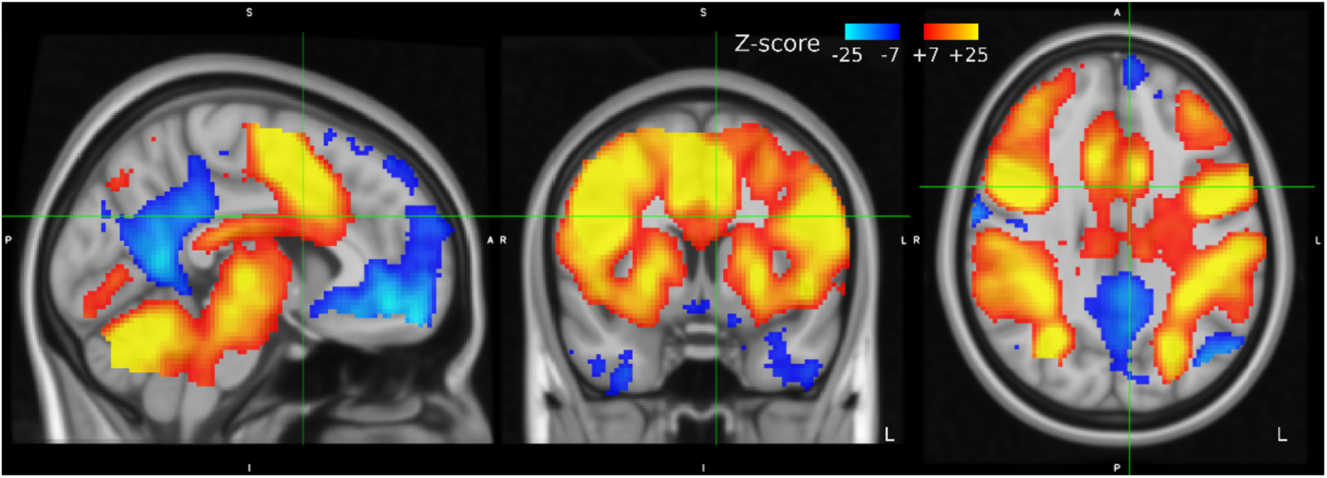
Group mean Z-statistics from the fMRI task, in MNI512 standard space; red-yellow positive, blue-cyan negative Z-scores on the range [7,25]

As for the comparison of BOLD responses assessed through fMRI with BOLD response assessed through fMRS (see Figure 6), there was a significant Spearman correlation between the strength of the BOLD response as assessed with fMRS and fMRI within the individually-prescribed VOI_fMRS,ACC_ (r=0.34, p_holm_=0.01, CI_95%_ [0.132,0.536]). A similar significant correlation was observed using a fixed VOI mask for all subjects, without accounting for individual placement (VOI_ACC_ r=0.33, p_holm_=0.01, CI_95%_ [0.11,0.52]). As can be seen from the correlation coefficients, both correlations were of moderate size. For control regions in the precuneus, occipital and temporal pole, no significant correlations were observed: r [CI_95%_] = 0.18 [−0.05, 0.39], 0.18 [−0.06, 0.39], 0.08 [−.16, .30] for VOI_precuneus_, VOI_occipital_ and VOI_temporal_ respectively, all p_unc._>0.1.

**Figure 6.**
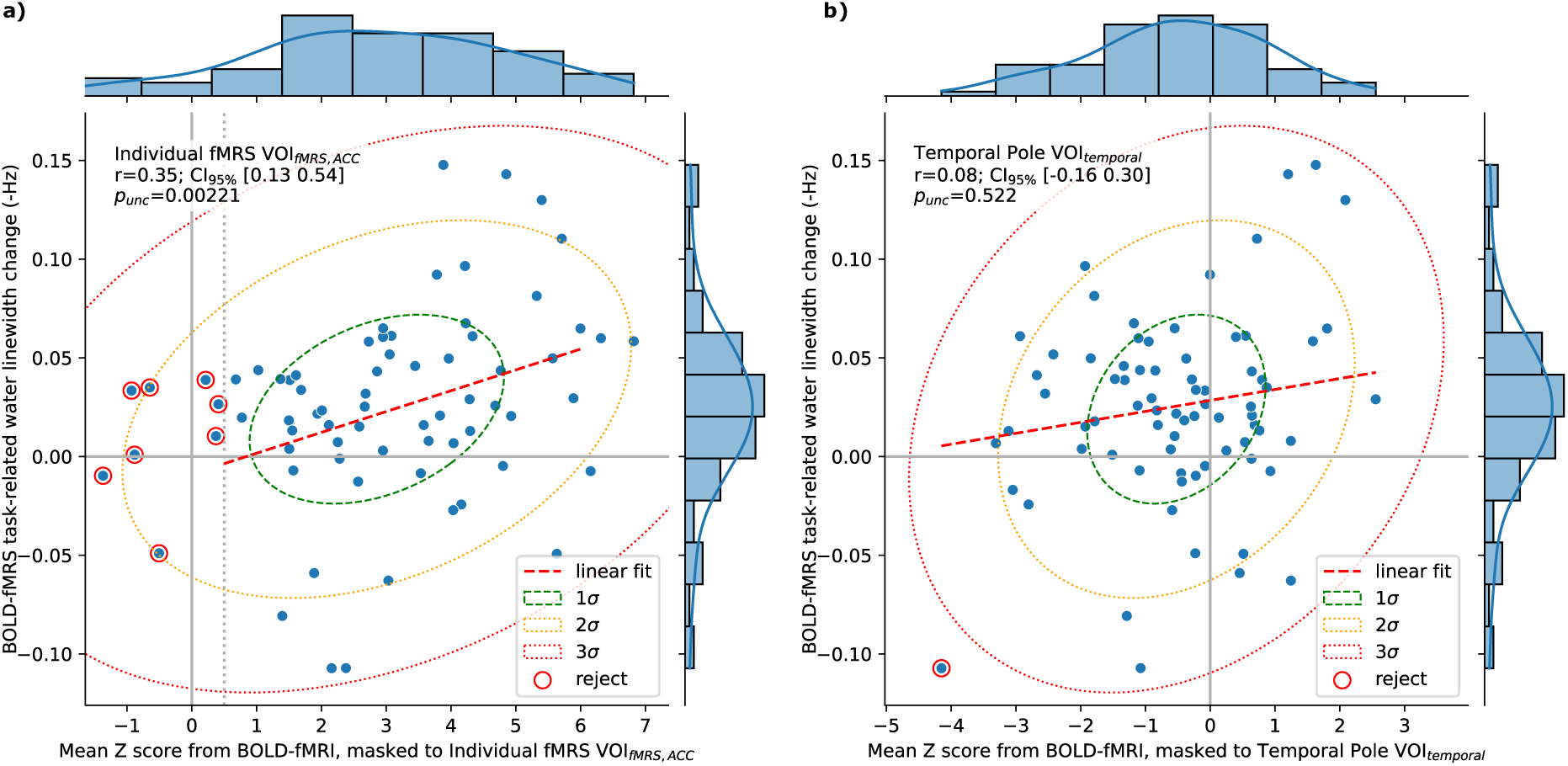
Relation of BOLD assessed by linewidth changes in the fMRS acquisition, to that observed in from the fMRI data regionally masked to (a) the individual fMRS voxel and (b) the temporal pole

#### 3.2.2 fMRS

From a total of 308 extracted spectra, rejection criterion R1 removed 25 fits (of which eight exhibited high NAA FWHM and 19 exhibited high GABA FWHM; some overlapped). Subsequently, one spectrum was removed by R2 (low NAA SNR) and eight extreme outliers were rejected by R3 (four for GABA+, five for Glx; some overlapped). Achieved spectral quality metrics (SNR and FWHM) and group average metabolite estimates after fitting and quality control are presented in Table 2; resultant spectra and mean fit for rest and active conditions are additionally presented in Figure 8. To verify the efficacy of linewidth matching, the standard deviation of Cho/Cr FWHM across extracted time bins (and rest) for each subject was evaluated before and after matching; median SD before matching 0.165 ± 0.05 Hz, reduced to 0.049 ± 0.021 Hz after matching: a significant improvement, p_holm_<0.001.

**Table 2.** Quality metrics and concentration estimates from the fMRS analysis, task-ON vs task-OFF

|  |  | Rest (task-OFF) | Active (task-ON) |
| --- | --- | --- | --- |
| SNR NAA | *** | $106.0 \pm 11.7$ | $44.7 \pm 5.24$ |
| FWHM Cho/Cr *before* linewidth matching | n.s. | $6.51 \pm 0.444$ | $6.44 \pm 0.453$ |
| FWHM Cho/Cr *after* linewidth matching | n.s. | $7.04 \pm 0.511$ | $7.04 \pm 0.512$ |
| FWHM NAA (Hz) | n.s. | $7.54 \pm 0.679$ | $7.43 \pm 0.642$ Hz |
| FWHM GABA+ (Hz) | n.s. | $17.8 \pm 1.95$ | $17.5 \pm 2.03$ |
| FWHM Glx (Hz) | n.s. | $12.7 \pm 1.26$ | $12.8 \pm 1.64$ |
| GABA+ (i.u., ≈mM) | n.s. | $2.93 \pm 0.457$ | $3.01 \pm 0.537$ |
| Glx (i.u., ≈mM) | ** | $14.8 \pm 1.48$ | $16.0 \pm 2.14$ |

Metabolite estimates separated by functional condition are shown in Figure 7; these results show a strongly significant increase in Glx following stimulus onsets (task-ON relative to task-OFF), of around 8.85% (p_holm_<0.001, CI_boot,95%_ [4.83, 13.9] %). While the figure may suggest a possible trend for GABA+, this was not significant (2.53%, p_unc._ =0.35, CI_boot,95%_ [−4.86,8.0]).

**Figure 7.**
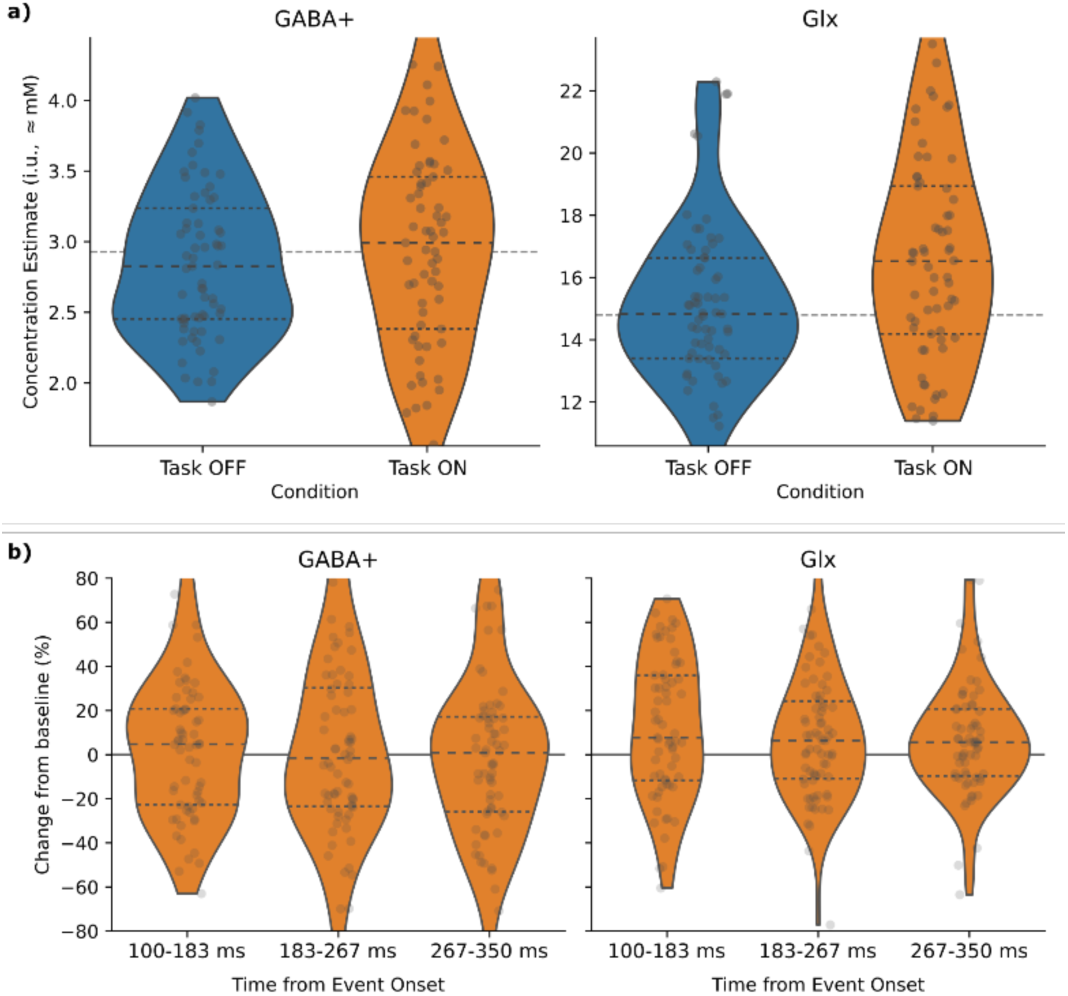
Metabolite concentration estimates for task-OFF vs task-ON conditions (a), and from spectra recorded at varying times after event onset, T_S-A_ (b)

Assessing the discrete T_S-A_ bins (Figure 7b), Glx appeared somewhat elevated in each time point, relative to task-OFF. This was statistically significant only for T_S-A_ = 100-183 ms (14.4% increase, p_holm_=0.02, CI_boot,95%_ [1.92,17.3]); elevation in other time bins was less pronounced and did not survive correction for multiple comparisons: 8.2% (p_unc._=0.03 CI_boot,95%_ [−3.66,11.2]) and 5.67% (p_unc._=0.02, CI_boot,95%_ [−2.45,9.74]) for T_S-A_ = 183-267 ms and 267-350 ms respectively. Pairwise comparison between time bins revealed no significant differences (all p_holm_>0.15; marginal p_unc._=0.049 between T_S-A_ = 100-183 and 267-350 ms), and no differences were seen for GABA+, either with respect to task-OFF or between time bins (all p_holm_>0.44). No significant differences were seen across portions of the task-ON block. Comparing metabolite estimates obtained for different task stimulus and response conditions (congruent vs incongruent, accurate and incongruent, inaccurate) also found no significant differences (all p_holm_>0.29).

### 3.3 Integrated Outcomes

Baseline metabolite concentration estimates (task-OFF) were assessed for correlation with fMRS BOLD estimates; only weak, non-significant correlations were obtained: r=-.18 (p_unc._=0.13, CI_95%_ [−0.398,0.054]) and r=-0.177 (p_unc._=0.135, CI_95%_ [−0.391,0.056]) for Glx and GABA+ respectively.

An exploratory analysis comparing behavioural measures (RT, RA, RA from incongruent trials only (RA_incong_) and incompatibility slowing) with fMRS-assessed BOLD and with metabolite estimates for GABA+ and Glx in task-OFF and task-ON periods revealed a significant correlation between incompatibility slowing and the strength of the BOLD response (r=0.39, p_holm_=0.04, CI_95%_ [0.151,0.586]). A possible negative correlation between RA_incong_ and task-OFF Glx levels (r=-0.379, p_holm_=0.13, p_unc._=0.006, CI_95%_ [−0.594, −0.112]) did not survive strict correction for multiple comparisons.

## 4 Discussion and Conclusions

### 4.1 Behavioural outcomes

The behavioural outcomes at group level showed strong incompatibility slowing effects (that is, the increase in reaction time for incongruent stimuli: RT_incongruent_-RT_congruent_) as is typical of Flanker tasks, and corresponding decreased response accuracy for incongruent stimuli. Accuracy effects remained significant also after normalising by reaction time (RA/RT). The degree of incompatibility slowing is consistent with existing studies (for example ^16–18,53^). Findings for absolute reaction times vary in the literature; reaction times obtained in the present study are consistent with some previous reports ^53^, but somewhat slower than reported elsewhere (often in the 300-400ms range). Error rates reported herein are consistent with those of Kopp et al. ^16^, although others have reported somewhat lower error rates, around 10% for the incongruent case. These variations are likely attributable to the geometry of the stimulus presentation (visual angle spanned by the flanker stimuli, as assessed in Kopp et al. ^16^), paradigm timing, the proportion of congruent vs incongruent stimuli and associated sequential dependencies.

For the more challenging incongruent stimuli, response accuracy was seen to improve between the fMRS and fMRI tasks, perhaps suggestive of learning effects which could be mitigated with a counter-balanced ordering (technical considerations notwithstanding, see section 4.2). Therefore, at least at group level, we conclude that the task was performed effectively and is expected to have elicited the desired cognitive load as evidenced in the BOLD fMRI outcomes.

### 4.2 BOLD functional outcomes

Group average BOLD contrast evaluated from the fMRI data exhibited a characteristic “task positive” structure consistent with a generalized extrinsic mode network ^54,55^ during task-ON blocks (see Figure 5). This included significant activation in the ACC region, as targeted by the fMRS voxel placement. Network nodes typically associated with the “resting state” default mode network ^56^ showed notably increased activation in the task-OFF blocks. In this way, we conclude that the implementation of the Flanker task used in this study resulted in the anticipated patterns of brain activation and deactivation in response to the presence or absence of task stimuli.

Although a significant correlation was seen between the BOLD signal assessed by the fMRI and fMRS methods, this correlation remained moderate. This could in part be attributed to the inherent variability of each measure, and intra-session variability in the BOLD response – although performing a similar task, these estimates were nonetheless derived from two discrete acquisitions. With fMRS acquired before fMRI, it could be that learning effects, fatigue, and other such factors mediate the strength of the BOLD signal in the later acquisition. Indeed, the behavioural outcomes suggest some learning effects. While a counter-balanced design may mitigate these factors, technical considerations relating to gradient heating and thermal drift ^9^ precluded counter-balancing in the current study.

In relation to behavioural outcomes, the significant correlation observed between fMRS-assessed BOLD and behavioural incompatibility slowing may be attributed to increased cognitive engagement required by those subjects demonstrating greater slowing.

### 4.3 GABA

A recent meta-analysis covering functional MRS studies of GABA and Glu/Glx ^57^ shows small effect sizes and large heterogeneity, both in experimental design and outcomes. Only one study in the meta-analysis investigated GABA changes during a cognitive (Stroop) task in the ACC, showing negative correlation between GABA change and BOLD signal change during task conditions ^58^. Outcomes across other tasks and locations (including ^59–65^) varied substantially, with no significant change found for GABA relative to baseline across studies: Hedge’s g −0.04 [−0.469, 0.386] n=11 and −0.01 [−0.25, 0.232] n=10 for studies reporting change in mean and percentage change respectively. In this context, our lack of significant results when examining GABA+ changes in relation to task is unremarkable.

Considering baseline GABA+ levels, an earlier meta-analysis ^66^ shows a number of studies reporting negative correlation between baseline GABA and the magnitude of BOLD measured during task performance ^67–71^. However, findings of positive associations (or non-associations) have also reported, suggesting the relation may be somewhat more nuanced ^72–75^. Although our results do not show a significant association, the obtained confidence interval [−0.391, 0.056] for Spearman correlation between baseline GABA and fMRS-assessed BOLD is not incompatible with documented negative correlations.

While there is no published consensus recommendation relating to the TR for GABA-edited MEGA-PRESS acquisitions, the TR parameter adopted in the present study was somewhat lower than commonly used (often around 2000 ms ^76,77^). As such, recovery of longitudinal magnetization will be somewhat reduced. Moreover, T_1_ weighting will be somewhat stronger, which may increase the contribution of macromolecule signals to the obtained edited spectrum due to their substantially shorter T_1_ ^78,79^. This trade-off allowed a higher number of functional events to be acquired, with a short ISI suitable for maintaining cognitive load. However, altered weighting and the potentially increased proportion of MM3co in the measured GABA+ signal may have limited the sensitivity to any more subtle GABA changes.

The present model and experimental design make no particular assumption as to the relative change in the edit-ON vs the edit-OFF sub-spectrum, in response to a functional task. Indeed, we may expect a change in MRS-visible GABA concentration to present more strongly in one of the sub-spectra. Explicit modelling of this, and tailoring the experimental design accordingly (targeting more transients from the more responsive sub-spectrum), in conjunction with a linear model which explicitly accounts for this, may yield stronger outcomes.

Finally, breaking the long fMRS acquisition into shorter segments, periodically adjusting the centre and editing frequencies and perhaps re-shimming to account for any changes over time (due to thermal drift, subject motion and so forth) may yield better editing outcomes than the single long acquisition, for future studies. Realtime frequency adjustment within a single acquisition may also mitigate the impact of frequency drift ^9^; in either case this factor would need to be incorporated into the model for spectral combination (section 2.4.2).

### 4.4 Glutamate and Glutamine (Glx) dynamics

The meta-analysis of Pasanta et al. ^57^ shows moderate effect sizes for Glu and Glx across studies, with block designs having somewhat tighter confidence intervals on effect size. The meta-analysis of Mullins ^80^ reports a mean change in Glutamate of 6.97% CI_95%_ [5.23, 8.72], although strongly dependent on both the experimental design (4.75% CI_95%_ [3.3, 6.2] for block designs, 13.43% [9.84, 17.02] for event-related designs) and the nature of the stimuli (2.318% [1.091, 3.545] for visual stimulus, 13.429% [9.839, 17.020] for pain). For the few studies reporting basic cognitive tasks in the ACC ^58,81,82^, typical changes were closer to 3%.

The changes reported in the present study (8.85% increase, CI_boot,95%_ [4.83, 13.9]) are comparatively strong; it is unlikely that a change of this magnitude, on the observed timescale, could be explained by metabolic processes (such as glutamate synthesis) alone. A more plausible explanation for such an increase could be a compartmental shift. It has been proposed that a substantial portion of Glu ^83^ and Gln ^84,85^ within the neuron may exist in pools where metabolite movement and tumbling is restricted – putatively including the synaptic vesicles. This restricted movement leads to faster T_2_ relaxation rates, and hence reduced visibility of the associated MRS signal, particularly at longer echo times such as used in the present study. During neural activity, glutamate released from vesicles may move to a compartment where it is more visible ^80,86^, such as the cytosol or synapse, leading to an increase in the apparent, measured signal.

As well as being the primary excitatory neurotransmitter, Glu plays a key role in normal energetic processes of neural cells, and in the synthesis of GABA ^87,88^. The proportion of Glu involved in each process is unclear, and current MRS techniques are generally not sufficiently selective to distinguish between the different compartments – although differing T_2_ across compartments means the relative visibility may be modulated by TE, with longer TEs being more sensitive to compartmental shifts ^80^. Disentangling these factors would serve to validate the hypothesized compartmental shift, and improve interpretability of the outcomes.

While the concentration estimates showed a robust increase in measured Glx between task-OFF and task-ON spectra, we note (with reference to Figure 8) that the 3.71 ppm sub-peak appears paradoxically slightly *reduced* in amplitude, while the most notable increase is seen in the leftmost part of the 3.79 ppm sub-peak, with an apparent broadening of the Glx peak in that area. While subtle variations in the peak shape may be consistent with a compartmental shift, one might expect a compartment of slower T_2_ (increased MRS visibility) to yield somewhat sharper peaks – this was not evident in the present data. Another possible explanation is that the apparent increase at the leftmost edge of the 3.75 ppm C2 peak may reflect an increased contribution of Glutamine to the observed signal, with the Glutamine C2 peak at 3.764 ppm being 0.017 ppm left of the Glutamate C2 peak at 3.747 ppm.

**Figure 8.**
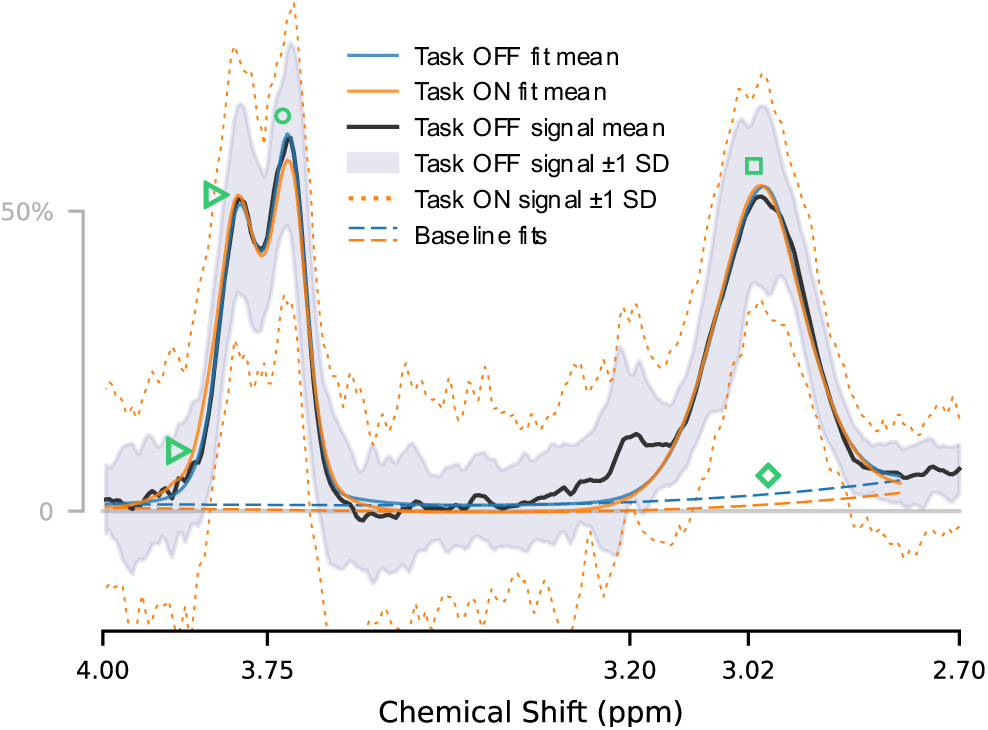
Mean extracted spectrum from the fMRS sequence for rest, showing ±1 SD range and mean fit for both rest and active conditions; note substantial overlap for GABA+ around 3.02 ppm □, slight differences in baseline fit around 3.02 ppm ◇ and in Glx shape at 3.71 ○ and 3.79 ppm ▷

The concentrations of glutamate and glutamine are closely linked, with the two in constant exchange in the neuronal-glial Glu-Gln cycle ^88,89^. While independent quantification of Glu and Gln may be feasible from GABA-edited data of sufficient quality ^90^, this approach has not been broadly adopted; indeed, there remains some question as to the reliability of Glx measurements obtained with from GABA-edited data ^91^. Given the comparatively low SNR of fMRS data, no attempt was made to separate Glu and Gln in the current data. Nonetheless, this may be feasible in further analysis using a basis-set fitting approach and strict acceptance criteria, or in future studies at higher field strengths and/or with dedicated sequences and judicious choice of sub-echo timings ^92^.

The typical trajectory of the glutamate response over time remains uncertain; while evidence from block-related designs suggests a gradual increase (around 16-20 seconds from the start of a block) likely relating to increased metabolism associated with neural activity, event-related studies suggest a separate process on a shorter timescale (returning to baseline within 3-4 seconds), related directly to neurotransmitter release from the vesicles ^19,80,93^. The event timing in the present study (T_S-A_ nominally 100 – 350 ms) was chosen to cover the range examined by Apšvalka et al. ^19^, as well as most of the evoked gamma-band power changes reported by Lally et al. ^93^. While statistically robust changes were only observed in the earlier time section, we do note a slight tendency towards baseline in subsequent times, as well as greater variance in the earlier part – perhaps suggestive of substantial inter-individual variability in the timing and onset of the initial response, which may normalise later in the response. Although the block timing of our study is well-suited to elicit a strong BOLD response and to assess the putative slower glutamate response function (GRF), with long task-OFF periods to allow for a robust return to baseline, the event timings (ISI ~1500 ms) were not optimal for unambiguous separation of adjacent events if we assume a return to baseline after 3-4 seconds for the faster GRF. Future studies primarily intending to model the GRF should ensure a longer interval between successive events, perhaps coupled with a longer TR for more optimal GABA measurement.

While the present analysis makes no particular assumptions regarding the shape of the GRF (beyond the restricted T_S-A_ range), we note that recent studies have begun to investigate putative metabolite response functions ^94^; once an appropriate response function is determined, incorporation of this into the present model (in place of the coarse binning) is trivially accomplished, and may well improve sensitivity to subtle variations. Furthermore, while the present model reconstructs discrete sub-spectra which are fit independently using existing 1D modelling tools, we note recent developments towards simultaneous, 2D fitting of multiple spectra linked by an arbitrary model, such as that implemented in the FSL-MRS dynamic fitting module ^95^. With appropriate constraints, incorporating our linear model into such an approach may further improve fitting performance, particularly for lower-SNR cases.

### 4.5 Conclusions

In the current study, we present a MEGA-PRESS sequence adaption suitable for the concurrent measurement of time-resolved GABA+, Glx and BOLD from a single voxel, at a ubiquitous 3T field strength. We additionally present a novel linear model for extracting spectra modelled from functional stimuli, building on well-established processing and quantification tools. With these tools, we demonstrate a robust increase in measured Glx concentration in the ACC in response to a task stimulus, measured concurrently with regional BOLD response. The latter is shown to significantly correlate with the BOLD response as assessed by traditional fMRI methods. Findings for Glx are consistent with theoretical models, and both fMRS and fMRI findings align with existing literature. Whilst identifying a number of readily achievable optimisations for future usage (particularly with regards to GABA sensitivity), we conclude that the acquisition and analysis methods documented herein are effective for the concurrent measurement of GABA, Glx and BOLD, in relation to a functional task.

## 5 Author Contributions

ARC: Conceptualization, Methodology, Software, Validation, Formal analysis, Investigation, Data Curation, Writing – Original Draft

GD: Conceptualization, Methodology, Validation, Investigation, Writing – Review & Editing

LE: Methodology, Validation, Resources, Writing – Review & Editing

KK: Data Curation, Writing – Review & Editing

RN: Methodology, Software, Writing – Review & Editing

LBS: Investigation, Data Curation, Writing – Review & Editing

EJ: Conceptualization, Resources, Writing – Review & Editing, Project administration

KH Conceptualization, Methodology, Resources, Writing – Review & Editing, Supervision, Project administration, Funding acquisition

## List of Abbreviations

(f)MRS: (functional) Magnetic Resonanse Spectroscopy
ACC: Anterior Cingulate Cortex
BOLD: Blood Oxygen Level Dependent
CHESS: CHEmically Selective Saturation pulses, used for water suppression
Cho: Choline
CI_(boot,)95%_: 95% confidence interval ((boot) denotes bootstrap CI)
Cr: Creatine
CSF: CerebroSpinal Fluid
DIFF: Difference spectrum (edit-ON – edit-OFF)
EPI: Echo-Planar Imaging
FFT: Fast Fourier Transform
fMRI: functional Magnetic Resonance Imaging
FSPGR: Fast SPoiled GRadient sequence (used for T_1_-weighted structural acquisition)
FWHM: Full Width at Half Maximum (linewidth)
GABA: γ-aminobutyric acid
GABA+: GABA plus underlying coedited (macromolecule) signal MM3co
Glu: Glutamate
GRF: Glutamate Response Function
HRF: Haemodynamic Response Function, characteristic of BOLD response
ISI: Interstimulus Interval
MAD: Median Absolute Deviation
MM3co: Coedited (macromolecule) signal around 3.0 ppm
NAA: N-acetylaspartate
p_holm_: Holm-Bonferroni adjusted p-value
PPG: Photoplethysmographic data (for pulse measurement)
p_unc_: Uncorrected p-value
R[1-3]: Rejection criteria (see 2.4.3)
RA_(cong/incong):_: Response Accuracy, in relation to the behavioural task (for congruent/incongruent stimuli)
rs-fMRI: resting state fMRI
RT: Reaction Time (in relation to behavioural task)
SNR: Signal-to-Noise Ratio
SOA: Stimulus Onset Asynchrony
SVD: Singular Value Decomposition
T_1_: Longitudinal relaxation time
T_2_*: Effective transverse relaxation time
TE: Echo Time
TPM: Tissue Probability Map
TR: Repetition Time
T_S-A_: Time from Stimulus to Acquisition
VOI_(region)_: Volume of Interest (in nominated region)
WREF: Water-unsuppressed reference transient

## 6 Acknowledgements

This study was funded by the European Research Council (ERC) grant #249516 and by the Western Norway Health Authorities (Helse-Vest) grant #912045 to Kenneth Hugdahl. We are grateful for the radiographers at Haukeland University Hospital: Roger Barndon, Christel Jansen, Turid Randa, Trond Øveraas, Eva Øksnes and Tor Erlend Fjørtoft, for their time and patience with data collection throughout this study.

## 8 Data availability statement

In accordance with data sharing regulations imposed by the Western Norway Ethical Committee (REK-Vest) (https://rekportalen.no/), data can be shared by request to the corresponding author, and after written permission from the REK-Vest.

Our custom MEGA-PRESS sequence is based on proprietary GE code; the authors are in principle willing to share details on our local adaptions through the appropriate vendor-facilitated channels.

## A MRSinMRS checklist

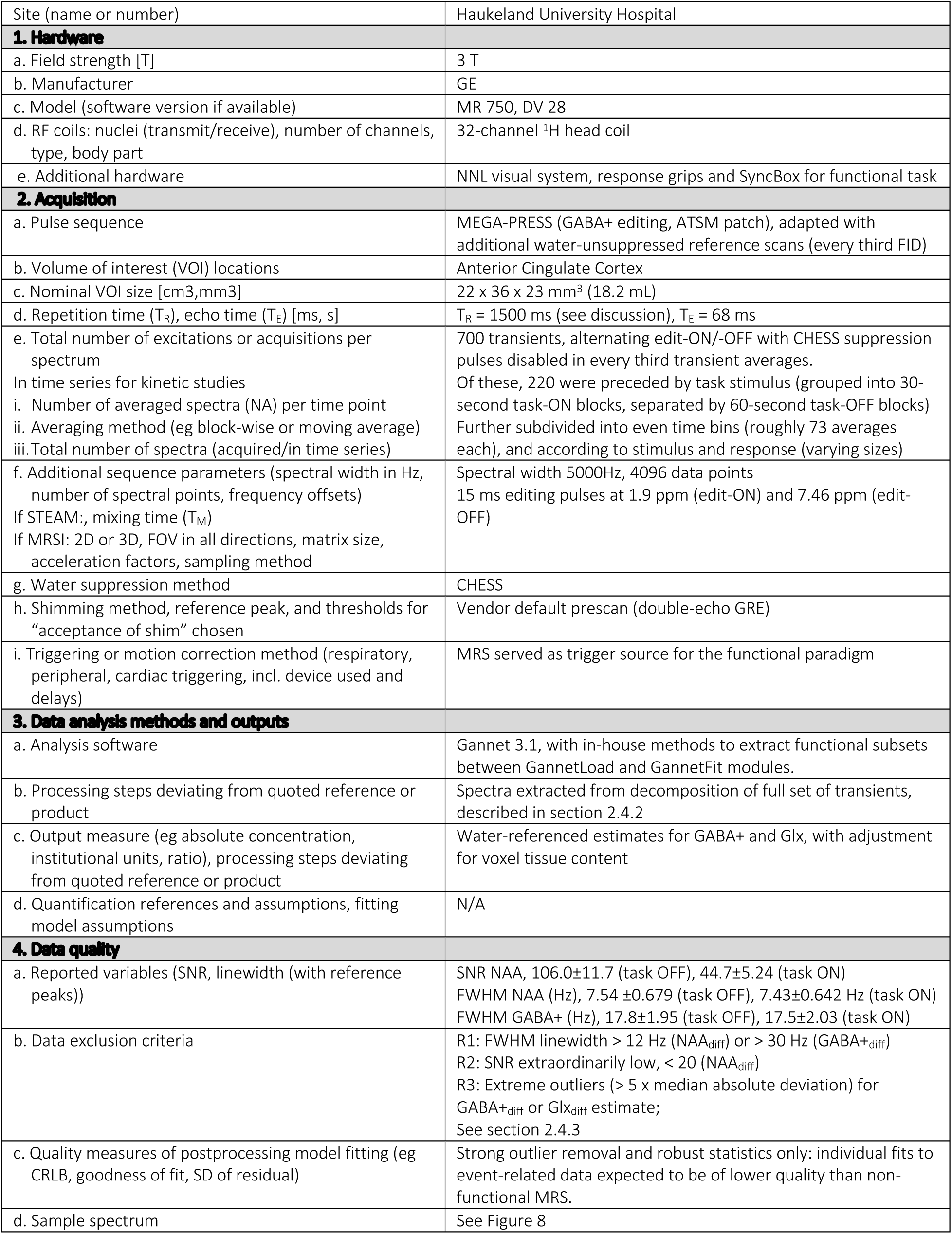

## B Supplementary Methods

**Supplementary Figure 1:**
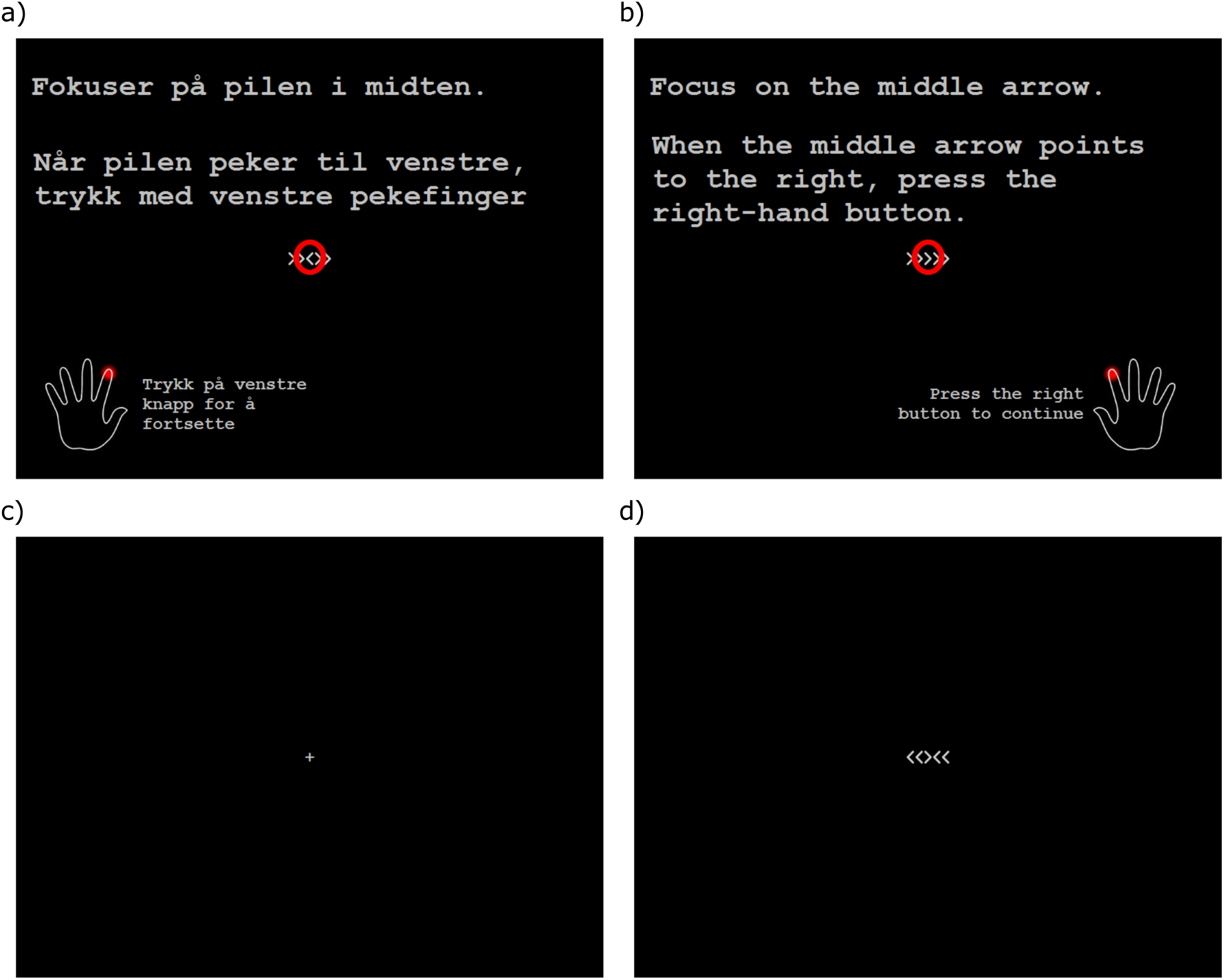
Sample instruction screens in Norwegian (a) and English (b); fixation (c) and sample Flanker stimulus (d)

## C BOLD fMRI outcome

Group average response from the BOLD fMRI task yielded an extensive cluster of activation during task-ON periods, spanning fronto-temporo-parietal regions and the ACC/SMA. The single huge cluster described in Supplementary Table 1 is characterised by numerous local maxima detailed in Supplementary Table 2. Task-OFF periods showed significant clusters in the parietal/precuneus and ventromedial inferior frontal regions, with local maxima detailed in Supplementary Table 3. Statistical Z-maps showing functional response are also presented in Supplementary Figure 2.

**Supplementary Table 1:** Major clusters identified from the BOLD-fMRI data, for Task ON and Task OFF periods; local maxima and associated structure are detailed in subsequent tables.

| Cluster Index | Num Voxels | P | -log<sub>10</sub>(P) | Z-MAX | X | Y | Z | COG X | COG Y | COG Z |
| --- | --- | --- | --- | --- | --- | --- | --- | --- | --- | --- |
| **Task ON** |  |  |  |  |  |  |  |  |  |  |
| 1 | 108966 | 0 | 661 | 40.1 | -4 | 4 | 48 | 2.08 | -23.3 | 14.8 |
| **Task OFF** |  |  |  |  |  |  |  |  |  |  |
| 9 | 23652 | 0 | 240 | 25 | -6 | 44 | -10 | -0.982 | 3.98 | 2.79 |
| 8 | 667 | 1.96E-24 | 23.7 | 17.8 | -44 | -72 | 28 | -46.4 | -66.3 | 24.6 |
| 7 | 314 | 1.00E-15 | 15 | 10.1 | 28 | 32 | 50 | 25.8 | 30.5 | 44.3 |
| 6 | 182 | 1.18E-11 | 10.9 | 9.94 | -60 | -8 | -18 | -56.2 | -9.97 | -16 |
| 5 | 182 | 1.18E-11 | 10.9 | 15.7 | -38 | -14 | 12 | -38.4 | -15.8 | 9.82 |
| 4 | 132 | 7.24E-10 | 9.14 | 9.7 | 20 | -30 | 58 | 20.4 | -30.2 | 61.7 |
| 3 | 118 | 2.51E-09 | 8.6 | 10.2 | 46 | -68 | 24 | 47.4 | -63.6 | 20.2 |
| 2 | 24 | 8.84E-05 | 4.05 | 7.77 | -60 | -4 | 10 | -59.5 | -3.24 | 8.67 |
| 1 | 10 | 0.00102 | 2.99 | 5.65 | 10 | 56 | 38 | 9.19 | 57 | 37.6 |

**Supplementary Figure 2:**
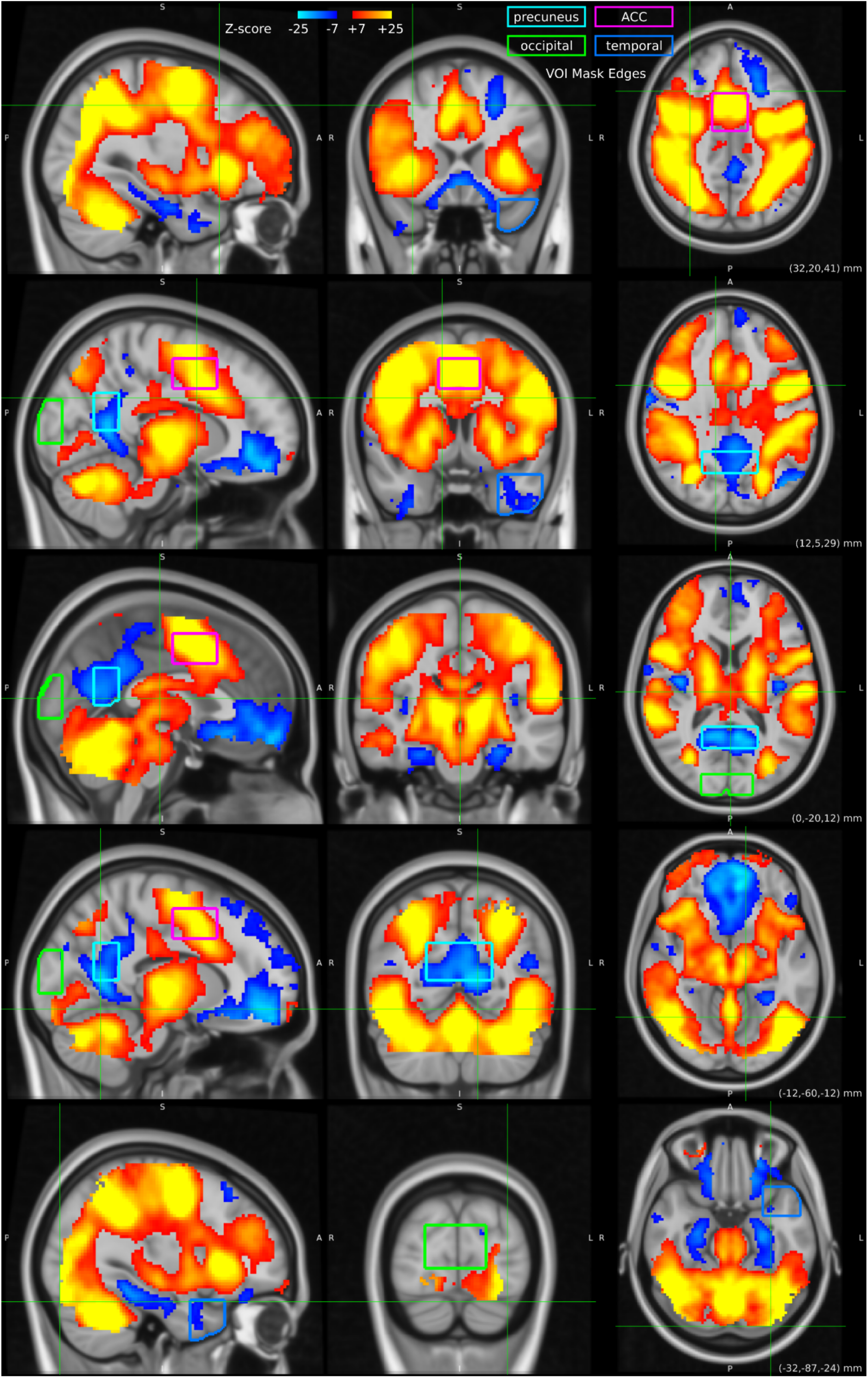
Z-statistic map from the BOLD-fMRI analysis, group level; VOI boundaries are also shown.

**Supplementary Table 2:** Local maxima from the **Task-ON** activation, with associated structures from the Harvard-Oxford Cortical and Subcortical Structural Atlases ^14,96–98^, and Cerebellar Atlas ^99^

| Cl# | Side | X | Y | Z | Atlas Structure 1, probability (%) | Atlas Structure 2, probability (%) | Atlas Structure 3, probability (%) |
| --- | --- | --- | --- | --- | --- | --- | --- |
| 1 | L | -4 | 4 | 48 | Juxtapositional Lobule Cortex (formerly Supplementary Motor Cortex) 62.0000 | Cingulate Gyrus, anterior division 15.0000 | Paracingulate Gyrus 10.0000 |
| 1 | L | -28 | -8 | 44 | Precentral Gyrus 23.0000 | Superior Frontal Gyrus 1.0000 | Middle Frontal Gyrus 1.0000 |
| 1 | L | -4 | 8 | 44 | Paracingulate Gyrus 38.0000 | Cingulate Gyrus, anterior division 30.0000 | Juxtapositional Lobule Cortex (formerly Supplementary Motor Cortex) 23.0000 |
| 1 | L | -50 | -2 | 36 | Precentral Gyrus 66.0000 | Middle Frontal Gyrus 2.0000 |  |
| 1 | R | 42 | -2 | 42 | Precentral Gyrus 43.0000 | Middle Frontal Gyrus 4.0000 |  |
| 1 | R | 4 | 6 | 48 | Juxtapositional Lobule Cortex (formerly Supplementary Motor Cortex) 50.0000 | Paracingulate Gyrus 24.0000 | Cingulate Gyrus, anterior division 13.0000 |
| 1 | L | -46 | -72 | -10 | Lateral Occipital Cortex, inferior division 67.0000 | Occipital Fusiform Gyrus 7.0000 | Inferior Temporal Gyrus, temporooccipital part 1.0000 |
| 1 | L | -44 | -4 | 42 | Precentral Gyrus 36.0000 | Middle Frontal Gyrus 4.0000 |  |
| 1 | R | 32 | -4 | 48 | Precentral Gyrus 23.0000 | Middle Frontal Gyrus 20.0000 | Superior Frontal Gyrus 4.0000 |
| 1 | R | 48 | 6 | 28 | Precentral Gyrus 49.0000 | Inferior Frontal Gyrus, pars opercularis 8.0000 |  |
| 1 | R | 32 | -46 | 36 | Superior Parietal Lobule 11.0000 | Angular Gyrus 7.0000 | Supramarginal Gyrus, posterior division 6.0000 |
| 1 | L | -30 | -56 | 44 | Superior Parietal Lobule 38.0000 | Angular Gyrus 10.0000 | Lateral Occipital Cortex, superior division 8.0000 |
| 1 | R | 8 | -70 | -26 | Right VI 68.0000 | Vermis VI 10.0000 | Right Crus I 1.0000 |
| 1 | L | -30 | -62 | -36 | Left Crus I 19.0000 | Left VI 2.0000 |  |
| 1 | L | -32 | -56 | -38 | Left Crus I 4.0000 |  |  |
| 1 | L | -42 | -38 | 34 | Supramarginal Gyrus, anterior division 6.0000 | Supramarginal Gyrus, posterior division 2.0000 | Angular Gyrus 1.0000 |
| 1 | - | 0 | -64 | -26 | Vermis VI 92.0000 |  |  |
| 1 | L | -6 | -72 | -26 | Left VI 57.0000 | Vermis VI 33.0000 | Left Crus I 4.0000 |
| 1 | R | 30 | -52 | 40 | Angular Gyrus 17.0000 | Superior Parietal Lobule 14.0000 | Lateral Occipital Cortex, superior division 1.0000 |
| 1 | R | 30 | -56 | -34 | Right VI 70.0000 | Right Crus I 14.0000 |  |
| 1 | R | 46 | -64 | -16 | Lateral Occipital Cortex, inferior division 39.0000 | Occipital Fusiform Gyrus 22.0000 | Inferior Temporal Gyrus, temporooccipital part 12.0000 |
| 1 | R | 12 | -14 | 0 | Right Thalamus 72.6343 | Right Cerebral White Matter 27.2491 |  |
| 1 | R | 30 | -56 | 46 | Superior Parietal Lobule 32.0000 | Lateral Occipital Cortex, superior division 19.0000 | Angular Gyrus 10.0000 |
| 1 | L | -32 | 20 | 2 | Insular Cortex 68.0000 | Frontal Orbital Cortex 1.0000 | Frontal Operculum Cortex 1.0000 |
| 1 | R | 28 | -64 | 34 | Lateral Occipital Cortex, superior division 51.0000 | Angular Gyrus 2.0000 |  |
| 1 | L | -12 | -18 | 2 | Left Thalamus 99.4672 | Left Cerebral White Matter 0.5328 |  |
| 1 | L | -42 | -14 | 52 | Precentral Gyrus 56.0000 | Postcentral Gyrus 12.0000 |  |
| 1 | R | 26 | -66 | 40 | Lateral Occipital Cortex, superior division 60.0000 | Precuneus Cortex 3.0000 |  |
| 1 | R | 32 | -70 | 18 | Lateral Occipital Cortex, superior division 14.0000 | Lateral Occipital Cortex, inferior division 1.0000 |  |
| 1 | L | -32 | -86 | -16 | Lateral Occipital Cortex, inferior division 29.0000 | Occipital Fusiform Gyrus 27.0000 | Occipital Pole 5.0000 |
| 1 | L | -28 | -72 | 20 | Lateral Occipital Cortex, superior division 17.0000 |  |  |
| 1 | L | -20 | -52 | -30 | Left VI 5.0000 |  |  |

**Supplementary Table 3.**
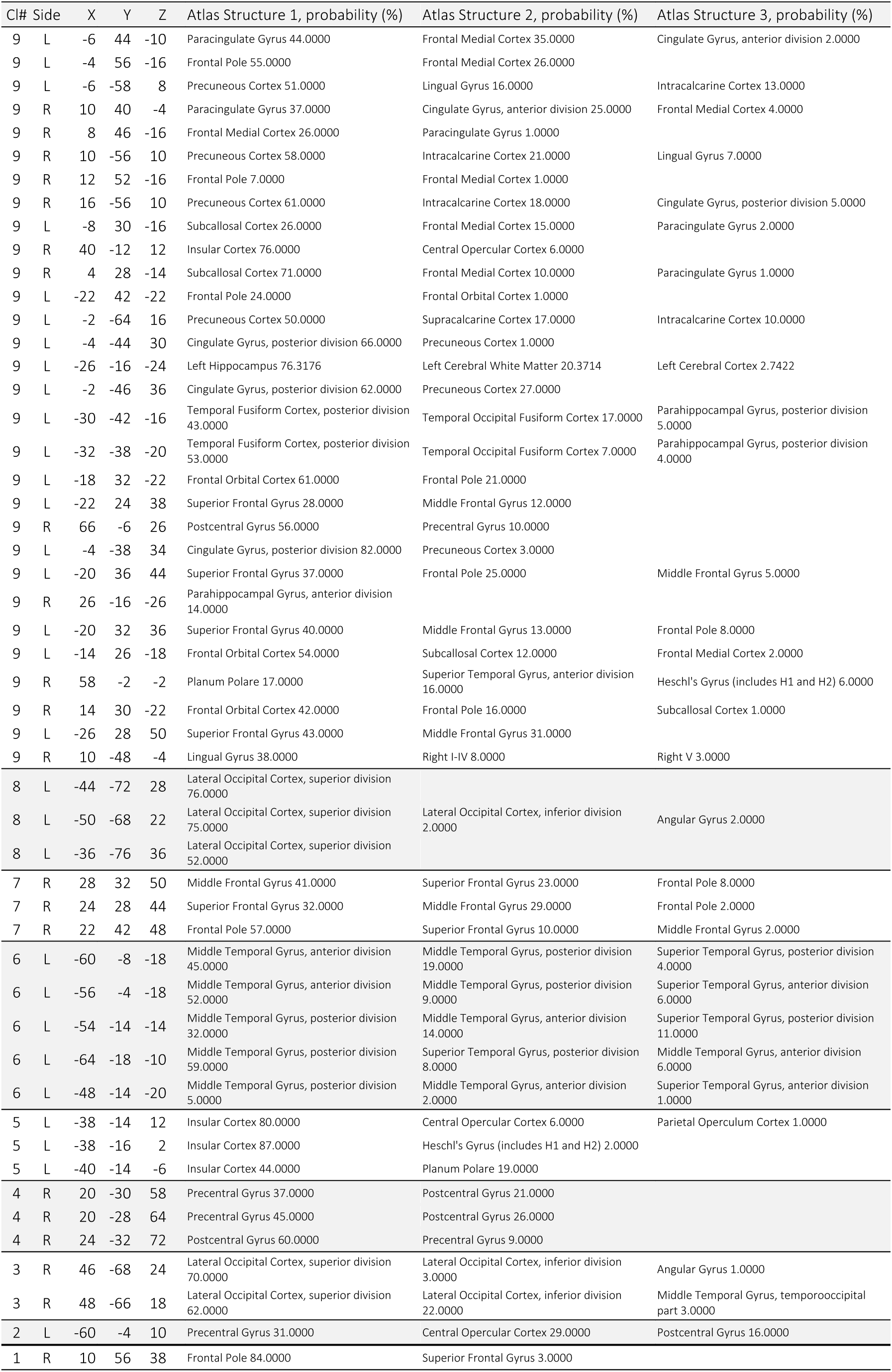
Local maxima from the **Task-OFF** activation, with associated structures from the Harvard-Oxford Cortical and Subcortical Structural Atlases ^14,96–98^, and Cerebellar Atlas ^99^

## References

1. Moonen CTW, Bandettini PA, Aguirre GK, eds. Functional MRI. Springer; 1999.

2. Ulmer S, Jansen O, eds. FMRI: Basics and Clinical Applications. Third edition. Springer; 2020.

3. Hugdahl K, Sommer IE. Auditory Verbal Hallucinations in Schizophrenia From a Levels of Explanation Perspective. Schizophr Bull. 2018;44(2):234–241. doi:10.1093/schbul/sbx142

4. Bertholdo D, Watcharakorn A, Castillo M. Brain Proton Magnetic Resonance Spectroscopy. Neuroimaging Clin N Am. 2013;23(3):359–380. doi:10.1016/j.nic.2012.10.002

5. Mescher M, Merkle H, Kirsch J, Garwood M, Gruetter R. Simultaneous in vivo spectral editing and water suppression. NMR Biomed. 1998;11(6):266–272. doi:10.1002/(sici)1099-1492(199810)11:6<266::aid-nbm530>3.0.co;2-j

6. Rothman DL, Petroff OA, Behar KL, Mattson RH. Localized 1H NMR measurements of gamma-aminobutyric acid in human brain in vivo. Proc Natl Acad Sci. 1993;90(12):5662–5666. doi:10.1073/pnas.90.12.5662

7. Near J, Simpson R, Cowen P, Jezzard P. Efficient γ-aminobutyric acid editing at 3T without macromolecule contamination: MEGA-SPECIAL. NMR Biomed. 2011;24(10):1277–1285. doi:10.1002/nbm.1688

8. Eriksen BA, Eriksen CW. Effects of noise letters upon the identification of a target letter in a nonsearch task. Percept Psychophys. 1974;16(1):143–149. doi:10.3758/BF03203267

9. Hui SCN, Mikkelsen M, Zöllner HJ, et al. Frequency drift in MR spectroscopy at 3T. NeuroImage. 2021;241:118430. doi:10.1016/j.neuroimage.2021.118430

10. Lin A, Andronesi O, Bogner W, et al. Minimum Reporting Standards for in vivo Magnetic Resonance Spectroscopy (MRSinMRS): Experts’ consensus recommendations. NMR Biomed. Published online February 9, 2021. doi:10.1002/nbm.4484

11. Ashburner J, Friston KJ. Unified segmentation. NeuroImage. 2005;26(3):839–851. doi:10.1016/j.neuroimage.2005.02.018

12. Blaiotta C, Freund P, Cardoso MJ, Ashburner J. Generative diffeomorphic atlas construction from brain and spinal cord MRI data. Published online July 5, 2017. Accessed January 30, 2023. http://arxiv.org/abs/1707.01342

13. Blaiotta C, Jorge Cardoso M, Ashburner J. Variational inference for medical image segmentation. Comput Vis Image Underst. 2016;151:14–28. doi:10.1016/j.cviu.2016.04.004

14. Desikan RS, Ségonne F, Fischl B, et al. An automated labeling system for subdividing the human cerebral cortex on MRI scans into gyral based regions of interest. NeuroImage. 2006;31(3):968–980. doi:10.1016/j.neuroimage.2006.01.021

15. Jones SH, Hemsley DR, Gray JA. Contextual Effects on Choice Reaction Time and Accuracy in Acute and Chronic Schizophrenics Impairment in Selective Attention or in the Influence of Prior Learning? Br J Psychiatry. 1991;159(3):415–421. doi:10.1192/bjp.159.3.415

16. Kopp B, Mattler U, Rist F. Selective attention and response competition in schizophrenic patients. Psychiatry Res. 1994;53(2):129–139. doi:10.1016/0165-1781(94)90104-X

17. Ridderinkhof KR, Wylie SA, van den Wildenberg WPM, Bashore TR, van der Molen MW. The arrow of time: Advancing insights into action control from the arrow version of the Eriksen flanker task. Atten Percept Psychophys. 2021;83(2):700–721. doi:10.3758/s13414-020-02167-z

18. Stoffels EJ, van der Molen MW. Effects of visual and auditory noise on visual choice reaction time in a continuous-flow paradigm. Percept Psychophys. 1988;44(1):7–14. doi:10.3758/BF03207468

19. Apšvalka D, Gadie A, Clemence M, Mullins PG. Event-related dynamics of glutamate and BOLD effects measured using functional magnetic resonance spectroscopy (fMRS) at 3 T in a repetition suppression paradigm. NeuroImage. 2015;118:292–300. doi:10.1016/j.neuroimage.2015.06.015

20. Edden RAE, Puts NAJ, Harris AD, Barker PB, Evans CJ. Gannet: A batch-processing tool for the quantitative analysis of gamma-aminobutyric acid-edited MR spectroscopy spectra: Gannet: GABA Analysis Toolkit. J Magn Reson Imaging. 2014;40(6):1445–1452. doi:10.1002/jmri.24478

21. Klose U. In vivo proton spectroscopy in presence of eddy currents. Magn Reson Med. 1990;14(1):26–30. doi:10.1002/mrm.1910140104

22. Jiru F. Introduction to post-processing techniques. Eur J Radiol. 2008;67(2):202–217. doi:10.1016/j.ejrad.2008.03.005

23. Mikkelsen M, Tapper S, Near J, Mostofsky SH, Puts NAJ, Edden RAE. Correcting frequency and phase offsets in MRS data using robust spectral registration. NMR Biomed. 2020;33(10). doi:10.1002/nbm.4368

24. Near J, Edden R, Evans CJ, Paquin R, Harris A, Jezzard P. Frequency and phase drift correction of magnetic resonance spectroscopy data by spectral registration in the time domain: MRS Drift Correction Using Spectral Registration. Magn Reson Med. 2015;73(1):44–50. doi:10.1002/mrm.25094

25. Gasparovic C, Song T, Devier D, et al. Use of tissue water as a concentration reference for proton spectroscopic imaging. Magn Reson Med. 2006;55(6):1219–1226. doi:10.1002/mrm.20901

26. Killick R, Fearnhead P, Eckley IA. Optimal Detection of Changepoints With a Linear Computational Cost. J Am Stat Assoc. 2012;107(500):1590–1598. doi:10.1080/01621459.2012.737745

27. Arlot S, Celisse A, Harchaoui Z. A kernel multiple change-point algorithm via model selection. J Mach Learn Res. 2019;20(162).

28. Garreau D, Arlot S. Consistent change-point detection with kernels. Published online June 29, 2017. Accessed February 20, 2023. http://arxiv.org/abs/1612.04740

29. Truong C, Oudre L, Vayatis N. Selective review of offline change point detection methods. Signal Process. 2020;167:107299. doi:10.1016/j.sigpro.2019.107299

30. Jenkinson M, Bannister P, Brady M, Smith S. Improved Optimization for the Robust and Accurate Linear Registration and Motion Correction of Brain Images. NeuroImage. 2002;17(2):825–841. doi:10.1006/nimg.2002.1132

31. Smith SM. Fast robust automated brain extraction. Hum Brain Mapp. 2002;17(3):143–155. doi:10.1002/hbm.10062

32. Jenkinson M, Smith S. A global optimisation method for robust affine registration of brain images. Med Image Anal. 2001;5(2):143–156. doi:10.1016/S1361-8415(01)00036-6

33. Andersson JL, Jenkinson M, Smith S. Non-linear optimisation FMRIB technical report TR07JA1. Practice. Published online 2007.

34. Andersson JL, Jenkinson M, Smith S, others. Non-linear registration, aka Spatial normalisation FMRIB technical report TR07JA2. FMRIB Anal Group Univ Oxf. 2007;2(1):e21.

35. Grabner G, Janke AL, Budge MM, Smith D, Pruessner J, Collins DL. Symmetric Atlasing and Model Based Segmentation: An Application to the Hippocampus in Older Adults. In: Larsen R, Nielsen M, Sporring J, eds. Medical Image Computing and Computer-Assisted Intervention – MICCAI 2006. Vol 4191. Lecture Notes in Computer Science. Springer Berlin Heidelberg; 2006:58–66. doi:10.1007/11866763_8

36. Woolrich MW, Ripley BD, Brady M, Smith SM. Temporal Autocorrelation in Univariate Linear Modeling of FMRI Data. NeuroImage. 2001;14(6):1370–1386. doi:10.1006/nimg.2001.0931

37. Worsley KJ. Statistical analysis of activation images. Ch 14. In: Jezzard P, Matthews PM, Smith SM, eds. Functional MRI: An Introduction to Methods.; 2001:251–270.

38. McKinney W. Data Structures for Statistical Computing in Python. In: ; 2010:56–61. doi:10.25080/Majora-92bf1922-00a

39. Harris CR, Millman KJ, van der Walt SJ, et al. Array programming with NumPy. Nature. 2020;585(7825):357–362. doi:10.1038/s41586-020-2649-2

40. SciPy 1.0 Contributors, Virtanen P, Gommers R, et al. SciPy 1.0: fundamental algorithms for scientific computing in Python. Nat Methods. 2020;17(3):261–272. doi:10.1038/s41592-019-0686-2

41. Vallat R. Pingouin: statistics in Python. J Open Source Softw. 2018;3(31):1026. doi:10.21105/joss.01026

42. Seabold S, Perktold J. statsmodels: Econometric and statistical modeling with python. In: ; 2010. https://www.statsmodels.org/

43. Hunter JD. Matplotlib: A 2D Graphics Environment. Comput Sci Eng. 2007;9(3):90–95. doi:10.1109/MCSE.2007.55

44. Waskom M. seaborn: statistical data visualization. J Open Source Softw. 2021;6(60):3021. doi:10.21105/joss.03021

45. Charlier F, Weber M, Izak D, et al. trevismd/statannotations: v0.5. Published online October 16, 2022. doi:10.5281/ZENODO.7213391

46. Fligner MA, Killeen TJ. Distribution-Free Two-Sample Tests for Scale. J Am Stat Assoc. 1976;71(353):210–213. doi:10.1080/01621459.1976.10481517

47. Shapiro SS, Wilk MB. An Analysis of Variance Test for Normality (Complete Samples). Biometrika. 1965;52(3/4):591. doi:10.2307/2333709

48. Wilcoxon F. Individual Comparisons by Ranking Methods. Biom Bull. 1945;1(6):80. doi:10.2307/3001968

49. Holm S. A Simple Sequentially Rejective Multiple Test Procedure. Scand J Stat. 1979;6(2):65–70.

50. Bonferroni CE. Il calcolo delle assicurazioni su gruppi di teste. Studi Onore Profr Salvatore Ortu Carboni. Published online 1935:13–60.

51. Pernet C, Wilcox R, Rousselet G. Robust Correlation Analyses: False Positive and Power Validation Using a New Open Source Matlab Toolbox. Front Psychol. 2013;3. doi:10.3389/fpsyg.2012.00606

52. Rousselet GA, Pernet CR. Improving standards in brain-behavior correlation analyses. Front Hum Neurosci. 2012;6. doi:10.3389/fnhum.2012.00119

53. Davelaar EJ, Stevens J. Sequential dependencies in the Eriksen flanker task: A direct comparison of two competing accounts. Psychon Bull Rev. 2009;16(1):121–126. doi:10.3758/PBR.16.1.121

54. Hugdahl K, Raichle ME, Mitra A, Specht K. On the existence of a generalized non-specific task-dependent network. Front Hum Neurosci. 2015;9. doi:10.3389/fnhum.2015.00430

55. Hugdahl K, Kazimierczak K, Beresniewicz J, et al. Dynamic up- and down-regulation of the default (DMN) and extrinsic (EMN) mode networks during alternating task-on and task-off periods. Hahn A, ed. PLOS ONE. 2019;14(9):e0218358. doi:10.1371/journal.pone.0218358

56. Raichle ME, MacLeod AM, Snyder AZ, Powers WJ, Gusnard DA, Shulman GL. A default mode of brain function. Proc Natl Acad Sci. 2001;98(2):676–682. doi:10.1073/pnas.98.2.676

57. Pasanta D, He JL, Ford T, Oeltzschner G, Lythgoe DJ, Puts NA. Functional MRS studies of GABA and glutamate/Glx – A systematic review and meta-analysis. Neurosci Biobehav Rev. 2023;144:104940. doi:10.1016/j.neubiorev.2022.104940

58. Kühn S, Schubert F, Mekle R, et al. Neurotransmitter changes during interference task in anterior cingulate cortex: evidence from fMRI-guided functional MRS at 3 T. Brain Struct Funct. 2016;221(5):2541–2551. doi:10.1007/s00429-015-1057-0

59. Bednařík P, Tkáč I, Giove F, et al. Neurochemical and BOLD Responses during Neuronal Activation Measured in the Human Visual Cortex at 7 Tesla. J Cereb Blood Flow Metab. 2015;35(4):601–610. doi:10.1038/jcbfm.2014.233

60. Bezalel V, Paz R, Tal A. Inhibitory and excitatory mechanisms in the human cingulate-cortex support reinforcement learning: A functional Proton Magnetic Resonance Spectroscopy study. NeuroImage. 2019;184:25–35. doi:10.1016/j.neuroimage.2018.09.016

61. Cleve M, Gussew A, Reichenbach JR. In vivo detection of acute pain-induced changes of GABA+ and Glx in the human brain by using functional 1H MEGA-PRESS MR spectroscopy. NeuroImage. 2015;105:67–75. doi:10.1016/j.neuroimage.2014.10.042

62. Dwyer GE, Craven AR, Bereśniewicz J, et al. Simultaneous Measurement of the BOLD Effect and Metabolic Changes in Response to Visual Stimulation Using the MEGA-PRESS Sequence at 3 T. Front Hum Neurosci. 2021;15:644079. doi:10.3389/fnhum.2021.644079

63. Kolasinski J, Hinson EL, Divanbeighi Zand AP, Rizov A, Emir UE, Stagg CJ. The dynamics of cortical GABA in human motor learning. J Physiol. 2019;597(1):271–282. doi:10.1113/JP276626

64. Michels L, Martin E, Klaver P, et al. Frontal GABA Levels Change during Working Memory. Koenig T, ed. PLoS ONE. 2012;7(4):e31933. doi:10.1371/journal.pone.0031933

65. Floyer-Lea A, Wylezinska M, Kincses T, Matthews PM. Rapid Modulation of GABA Concentration in Human Sensorimotor Cortex During Motor Learning. J Neurophysiol. 2006;95(3):1639–1644. doi:10.1152/jn.00346.2005

66. Duncan NW, Wiebking C, Northoff G. Associations of regional GABA and glutamate with intrinsic and extrinsic neural activity in humans—A review of multimodal imaging studies. Neurosci Biobehav Rev. 2014;47:36–52. doi:10.1016/j.neubiorev.2014.07.016

67. Donahue MJ, Near J, Blicher JU, Jezzard P. Baseline GABA concentration and fMRI response. NeuroImage. 2010;53(2):392–398. doi:10.1016/j.neuroimage.2010.07.017

68. Hayes DJ, Duncan NW, Wiebking C, et al. GABAA Receptors Predict Aversion-Related Brain Responses: An fMRI-PET Investigation in Healthy Humans. Neuropsychopharmacology. 2013;38(8):1438–1450. doi:10.1038/npp.2013.40

69. Muthukumaraswamy SD, Evans CJ, Edden RAE, Wise RG, Singh KD. Individual variability in the shape and amplitude of the BOLD-HRF correlates with endogenous GABAergic inhibition. Hum Brain Mapp. 2012;33(2):455–465. doi:10.1002/hbm.21223

70. Muthukumaraswamy SD, Edden RAE, Jones DK, Swettenham JB, Singh KD. Resting GABA concentration predicts peak gamma frequency and fMRI amplitude in response to visual stimulation in humans. Proc Natl Acad Sci. 2009;106(20):8356–8361. doi:10.1073/pnas.0900728106

71. Northoff G, Walter M, Schulte RF, et al. GABA concentrations in the human anterior cingulate cortex predict negative BOLD responses in fMRI. Nat Neurosci. 2007;10(12):1515–1517. doi:10.1038/nn2001

72. Qin P, Duncan NW, Wiebking C, et al. GABAA receptors in visual and auditory cortex and neural activity changes during basic visual stimulation. Front Hum Neurosci. 2012;6. doi:10.3389/fnhum.2012.00337

73. Wiebking C, Duncan NW, Tiret B, et al. GABA in the insula — a predictor of the neural response to interoceptive awareness. NeuroImage. 2014;86:10–18. doi:10.1016/j.neuroimage.2013.04.042

74. Wiebking C, Duncan NW, Qin P, et al. External awareness and GABA-A multimodal imaging study combining fMRI and [^18^ F]flumazenil-PET: GABA and External Awareness. Hum Brain Mapp. 2014;35(1):173–184. doi:10.1002/hbm.22166

75. Jocham G, Hunt LT, Near J, Behrens TEJ. A mechanism for value-guided choice based on the excitation-inhibition balance in prefrontal cortex. Nat Neurosci. 2012;15(7):960–961. doi:10.1038/nn.3140

76. Deelchand DK, Marjańska M, Henry P, Terpstra M. MEGA-PRESS of GABA+: Influences of acquisition parameters. NMR Biomed. 2021;34(5). doi:10.1002/nbm.4199

77. Peek AL, Rebbeck TJ, Leaver AM, et al. A comprehensive guide to MEGA-PRESS for GABA measurement. Anal Biochem. 2023;669:115113. doi:10.1016/j.ab.2023.115113

78. Bhagwagar Z, Wylezinska M, Jezzard P, et al. Reduction in Occipital Cortex γ-Aminobutyric Acid Concentrations in Medication-Free Recovered Unipolar Depressed and Bipolar Subjects. Biol Psychiatry. 2007;61(6):806–812. doi:10.1016/j.biopsych.2006.08.048

79. Mullins PG, McGonigle DJ, O’Gorman RL, et al. Current practice in the use of MEGA-PRESS spectroscopy for the detection of GABA. NeuroImage. 2014;86:43–52. doi:10.1016/j.neuroimage.2012.12.004

80. Mullins PG. Towards a theory of functional magnetic resonance spectroscopy (fMRS): A meta-analysis and discussion of using MRS to measure changes in neurotransmitters in real time. Scand J Psychol. 2018;59(1):91–103. doi:10.1111/sjop.12411

81. Taylor R, Schaefer B, Densmore M, et al. Increased glutamate levels observed upon functional activation in the anterior cingulate cortex using the Stroop Task and functional spectroscopy. NeuroReport. 2015;26(3):107–112. doi:10.1097/WNR.0000000000000309

82. Taylor R, Neufeld RWJ, Schaefer B, et al. Functional magnetic resonance spectroscopy of glutamate in schizophrenia and major depressive disorder: anterior cingulate activity during a color-word Stroop task. Npj Schizophr. 2015;1(1):15028. doi:10.1038/npjschz.2015.28

83. Kauppinen RA, Pirttilä TRM, Auriola SOK, Williams SR. Compartmentation of cerebral glutamate *in situ* as detected by 1H/13C n.m.r. Biochem J. 1994;298(1):121–127. doi:10.1042/bj2980121

84. Rae C, Hare N, Bubb WA, et al. Inhibition of glutamine transport depletes glutamate and GABA neurotransmitter pools: further evidence for metabolic compartmentation: Role of glutamine transport in CNS metabolism. J Neurochem. 2003;85(2):503–514. doi:10.1046/j.1471-4159.2003.01713.x

85. Hancu I, Port J. The case of the missing glutamine. NMR Biomed. 2011;24(5):529–535. doi:10.1002/nbm.1620

86. Lea-Carnall CA, El-Deredy W, Stagg CJ, Williams SR, Trujillo-Barreto NJ. A mean-field model of glutamate and GABA synaptic dynamics for functional MRS. NeuroImage. Published online December 2022:119813. doi:10.1016/j.neuroimage.2022.119813

87. Mangia S, Giove F, Tkáč I, et al. Metabolic and Hemodynamic Events after Changes in Neuronal Activity: Current Hypotheses, Theoretical Predictions and *in vivo* NMR Experimental Findings. J Cereb Blood Flow Metab. 2009;29(3):441–463. doi:10.1038/jcbfm.2008.134

88. Rae CD. A Guide to the Metabolic Pathways and Function of Metabolites Observed in Human Brain 1H Magnetic Resonance Spectra. Neurochem Res. 2014;39(1):1–36. doi:10.1007/s11064-013-1199-5

89. Ramadan S, Lin A, Stanwell P. Glutamate and glutamine: a review of *in vivo* MRS in the human brain. NMR Biomed. 2013;26(12):1630–1646. doi:10.1002/nbm.3045

90. Sanaei Nezhad F, Anton A, Michou E, Jung J, Parkes LM, Williams SR. Quantification of GABA, glutamate and glutamine in a single measurement at 3 T using GABA-edited MEGA-PRESS. NMR Biomed. 2018;31(1). doi:10.1002/nbm.3847

91. Bell T, Boudes ES, Loo RS, et al. In vivo Glx and Glu measurements from GABA-edited MRS at 3 T. NMR Biomed. Published online January 28, 2020:e4245. doi:10.1002/nbm.4245

92. Snyder J, Wilman A. Field strength dependence of PRESS timings for simultaneous detection of glutamate and glutamine from 1.5 to 7T. J Magn Reson. 2010;203(1):66–72. doi:10.1016/j.jmr.2009.12.002

93. Lally N, Mullins PG, Roberts MV, Price D, Gruber T, Haenschel C. Glutamatergic correlates of gamma-band oscillatory activity during cognition: A concurrent ER-MRS and EEG study. NeuroImage. 2014;85:823–833. doi:10.1016/j.neuroimage.2013.07.049

94. Schrantee A, Berrington A, Pouwels PJ, Coolen BF, Gurney-Champion O, Najac CF. The effect of the glutamate response function on detection sensitivity for functional MRS: a simulation study. In: Proc. Intl. Soc. Mag. Reson. Med.; 2022. https://archive.ismrm.org/2022/1077.html

95. Clarke WT, Ligneul C, Cottaar M, Jbabdi S. Dynamic fitting of functional MRS, diffusion weighted MRS, and edited MRS using a single interface. In: Proc. Intl. Soc. Mag. Reson. Med. Vol 30. ; 2022:3115. https://archive.ismrm.org/2022/0309.html

96. Frazier JA, Chiu S, Breeze JL, et al. Structural Brain Magnetic Resonance Imaging of Limbic and Thalamic Volumes in Pediatric Bipolar Disorder. Am J Psychiatry. 2005;162(7):1256–1265. doi:10.1176/appi.ajp.162.7.1256

97. Goldstein JM, Seidman LJ, Makris N, et al. Hypothalamic Abnormalities in Schizophrenia: Sex Effects and Genetic Vulnerability. Biol Psychiatry. 2007;61(8):935–945. doi:10.1016/j.biopsych.2006.06.027

98. Makris N, Goldstein JM, Kennedy D, et al. Decreased volume of left and total anterior insular lobule in schizophrenia. Schizophr Res. 2006;83(2-3):155–171. doi:10.1016/j.schres.2005.11.020

99. Diedrichsen J, Balsters JH, Flavell J, Cussans E, Ramnani N. A probabilistic MR atlas of the human cerebellum. NeuroImage. 2009;46(1):39–46. doi:10.1016/j.neuroimage.2009.01.045

